# Nuclear stabilisation of p53 requires a functional nucleolar surveillance pathway

**DOI:** 10.1101/2021.01.21.427535

**Authors:** Katherine M. Hannan, Priscilla Soo, Mei S. Wong, Justine K. Lee, Nadine Hein, Maurits Evers, Kira D. Wysoke, Tobias D. Williams, Christian Montellese, Lorey K. Smith, Sheren J. Al-Obaidi, Lorena Núñez-Villacís, Perlita Poh, Megan Pavy, Jin-Shu He, Kate M. Parsons, Jeannine Diesch, Gaetan Burgio, Rita Ferreira, Zhi-Ping Feng, Cathryn M. Gould, Piyush B. Madhamshettiwar, Johan Flygare, Thomas J. Gonda, Kaylene J. Simpson, Ulrike Kutay, Richard B. Pearson, Christoph Engel, Nicholas J. Watkins, Ross D. Hannan, Amee J. George

**Affiliations:** ACRF Department of Cancer Biology and Therapeutics, John Curtin School of Medical Research, Australian National University, Acton 2601 Australia; Oncogenic Signalling and Growth Control Program, Peter MacCallum Cancer Centre, Melbourne 3000, Australia; Department of Biochemistry and Molecular Biology, University of Melbourne, Parkville 3010 Australia; Sir Peter MacCallum Department of Oncology, University of Melbourne, Parkville 3010 Australia; Institute for Cell and Molecular Biosciences, Newcastle University, Newcastle-Upon-Tyne, UK; Department of Biology, Institute of Biochemistry, ETH Zurich, Zurich, Switzerland; ANU Centre for Therapeutic Discovery, John Curtin School of Medical Research, Australian National University, Acton 2601 Australia; The John Curtin School of Medical Research, Australian National University, Acton 2601 Australia; Victorian Centre for Functional Genomics, Peter MacCallum Cancer Centre, Melbourne 3000, Australia; Lund Stem Cell Center, Lund University, BMC A12, Lund, Sweden; School of Pharmacy, University of Queensland, Brisbane 4102 Australia; Department of Biochemistry and Molecular Biology, Monash University, Clayton, 3800, Australia; Regensburg Center for Biochemistry, University of Regensburg, 93053 Regensburg, Germany; School of Biomedical Sciences, University of Queensland, St Lucia 4067, Australia; Department of Clinical Pathology, University of Melbourne, Parkville 3010 Australia

**Keywords:** Nucleolar surveillance pathway, nucleolus, p53, ribosome biogenesis, high-throughput screening, ribosomal proteins, stress

## Abstract

The nucleolar surveillance pathway (NSP) monitors nucleolar fidelity and responds to nucleolar stresses (i.e., inactivation of ribosome biogenesis) by mediating the inhibitory binding of ribosomal proteins (RPs) to mouse double minute 2 homolog (MDM2), a nuclear-localised E3 ubiquitin ligase, which results in p53 accumulation. Inappropriate activation of the NSP has been implicated in the pathogenesis of collection of human diseases termed “ribosomopathies”, while drugs that selectively activate the NSP are now in trials for cancer. Despite the clinical significance, the precise molecular mechanism(s) regulating the NSP remain poorly understood. Using genome-wide loss of function screens, we demonstrate the ribosome biogenesis (RiBi) axis as the most potent class of genes whose disruption stabilises p53. Furthermore, we identified a novel suite of genes critical for the NSP, including a novel mammalian protein implicated in 5S ribonucleoprotein particle (5S-RNP) biogenesis, HEATR3. By selectively disabling the NSP, we unexpectedly demonstrate that a functional NSP is required for the ability of all nuclear acting stresses tested, including DNA damage, to robustly induce p53 accumulation. Together, our data demonstrates that the NSP has evolved as the dominant central integrator of stresses that regulate nuclear p53 abundance, thus ensuring RiBi is hardwired to cellular proliferative capacity.

## Main

Mutations in the potent tumour suppressor protein p53 and its effector pathways occur in the majority of human cancers, and are therefore the subject of intense investigation. A key mechanism by which p53 is regulated is at the level of protein stabilisation, through the MDM2 protein, which induces ubiquitination, and subsequently proteasomal degradation of p53. DNA damage from ionising radiation or certain chemotherapeutic agents lead to the amino-terminal phosphorylation of p53, which prevents MDM2 binding, and results in p53 stabilisation. This triggers a number of anti-proliferative programs by activating or repressing key effector genes in a context-dependent manner^1^. The p53-MDM2 interaction is also antagonised by the tumour suppressor p14^ARF^ in response to oncogenic challenges^2^. More recently, a third mechanism of p53 stabilisation has been identified; the NSP, which is activated by acute disruptions to RiBi, resulting in inhibitory binding of certain RPs to MDM2, thus leading to increased abundance of nuclear p53 protein^3-5^. In contrast to the former, the precise mechanisms underlying p53 stabilisation in response to the NSP are poorly understood. For example, the ribosomal proteins RPL5 and RPL11 have been implicated as the central regulators of the NSP through their participation in the 5S-RNP complex that binds to and inactivates MDM2 in response to nucleolar stress^4,5^. However, other RP and non-RP genes have also been implicated in regulating the NSP signalling process, suggesting the definitive mechanism is yet to be resolved. It is also unclear why loss or inactivation of only certain ribosome-associated genes give rise to increased p53 stabilisation or are connected with ribosomopathies. Finally, the functional relationship of the NSP to the mechanisms underlying p53 stabilisation observed in response to ‘classic’ non-nucleolar stress pathways, such as proteasomal stress, hypoxia or DNA damage, is not clear.

To address these questions, we first identified the entire repertoire of genes whose deletion activates stress pathways leading to stabilisation of p53 in A549 (human lung adenocarcinoma, p53 wild-type) cells, by undertaking a high-throughput genome-wide RNA interference (RNAi) imaging-based screen measuring nuclear p53 accumulation using immunofluorescence (‘p53 stabilisation screen’, **Fig. 1a, Supplementary File 1**). The screen ‘cut-off’ was functionally defined as the minimum amount of p53 accumulation required to induce a significant cell-cycle defect **(Supplementary Fig. 1a-d**), which we identified as ∼2-fold increase in p53 protein expression. Applying this cut-off (log_2_ ≥ 1) to the screening dataset, 827 genes fulfilled this criterion (defined as ‘p53 positive’, **Fig. 1b**). We further interrogated the ‘p53 positive’ candidates to identify which molecular pathways/functions were enriched in the dataset using the KEGG network enrichment analysis feature of STRING^6^ (**Fig. 1c**, annotated version in **Supplementary Fig. 2a & Supplementary File 2**). This revealed an enrichment of six major classes of genes including: ribosome, nucleolus, proteasome, RNA splicing, cell cycle and RNA Polymerase II (Pol II). These classes were also broadly confirmed by gene ontology (GO) analysis (**Fig. 1d, Supplementary File 3**) and gene set enrichment analysis (GSEA, **Supplementary Fig. 2b**), resulting in GOs relating to RNP complex and RiBi, ribosomal RNA (rRNA) processing and rRNA metabolic processes being amongst the most significantly enriched. Moreover, intersecting our ‘p53 positive’ candidate list with the LOCATE subcellular localisation database^7^, we identified a significant over-representation of proteins localised to the nucleolus, nucleus and centrosome, and conversely, an under-representation of proteins located within the plasma membrane (**Fig. 1e, Supplementary File 4**). Collectively, these observations strongly support the notion that perturbations in RiBi and/or the nucleolus are a major, if not the most potent regulators of p53 accumulation.

**Figure 1:**
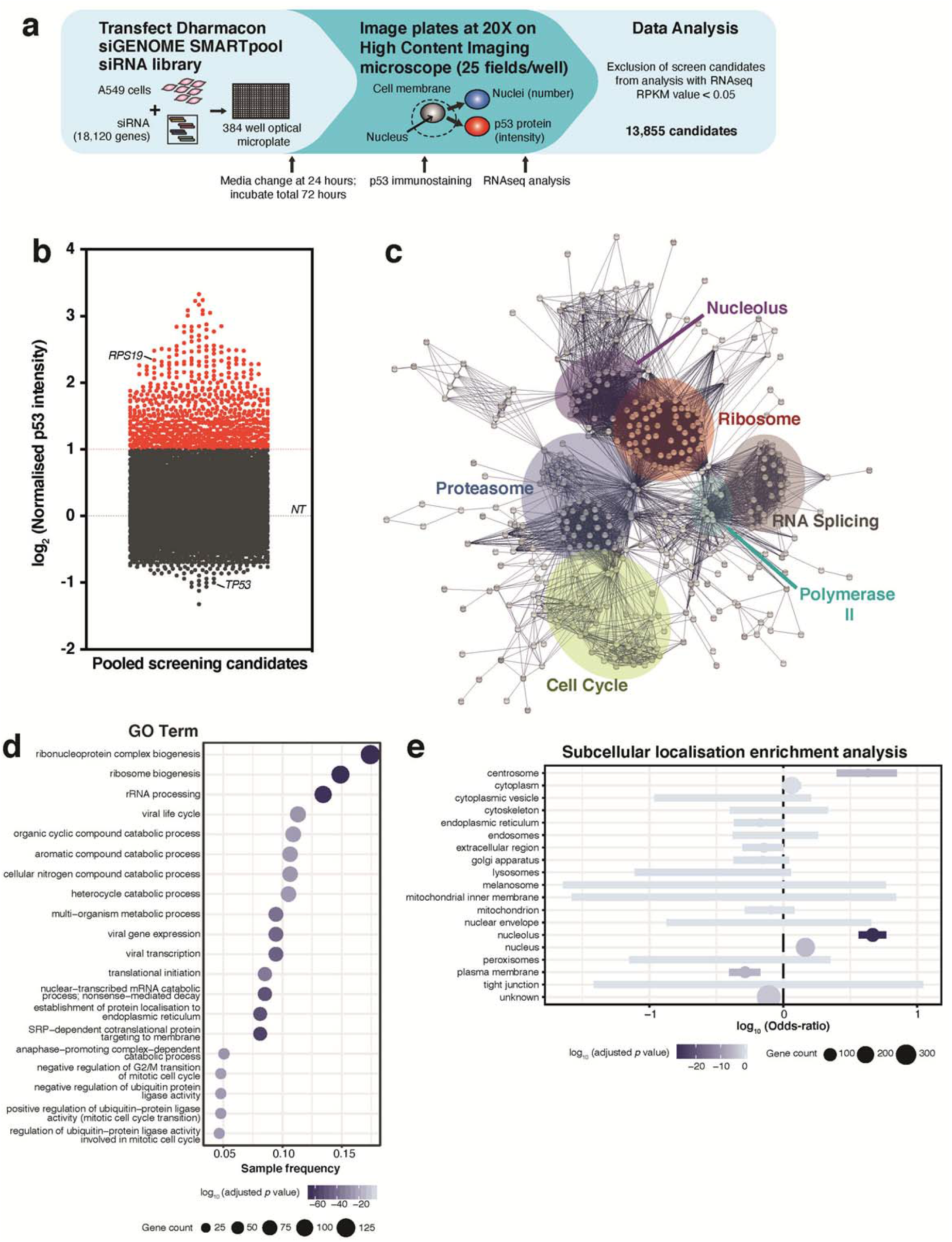
A genome-wide high-throughput screen reveals ribosome biogenesis and the nucleolus as central components for modulating p53 stabilisation. Schematic of the genome-wide high throughput screening approach in A549 (p53 wild-type) cells (**a**). A549 cells were transfected with the genome-wide Dharmacon siGENOME SMARTpool siRNA library for 72 hours, then nuclear p53 intensity and cell number measured using an immunofluorescence-based high-content (microscopy) imaging approach, with data normalised to non-targeting siRNA (NT) transfected cells (‘p53 stabilisation screen’). After intersection with RNAseq data from NT-transfected cells (a cut-off of reads per kilobase per million, RPKM, of 0.05 or greater, to ensure that candidates analysed are expressed in these cells), we determined the ‘expressed’ screening candidates to be 13,855. The ‘expressed’ candidates are graphed in (**b**); the top candidates (coloured in red) are those which were 2-fold or greater (log_2_ = 1); 827 candidates in total. The top candidates were then subjected to STRING (protein-protein interaction database) analysis using the KEGG network enrichment analysis feature, and visualised in Cytoscape (described in Methods, **c**) to identify clusters of similar proteins in the dataset (note: a fully annotated version of this figure is located in Supplementary Figure 2A). Gene ontology (GO) analysis of the top candidates was then performed, and simplified for graphical representation of the ‘Biological Process’ data using ClusterProfiler (**d**, approach outlined in Methods), which depicts the gene ontologies versus the gene ratio (ratio of number of query genes in the GO term and the total number of query genes). Subcellular localisation enrichment analysis of the top candidates from the screen (using the LOCATE database, **e**) was also performed; the log_10_ odds-ratio (OR) reflects the amount of enrichment/depletion (<0 indicates under-representation, >0 indicates over-representation of query genes in the corresponding category). The coloured bars represent 95% confidence intervals.

We initially focussed specifically on RP genes given their prominence in the dataset; ∼80% of the RPs screened were ‘p53 positive’ when depleted, including RP genes associated with Diamond-Blackfan Anaemia (DBA; e.g. *RPS19, RPL35A, RPS7, RPS10, RPS24, RPS26, RPL26*)^8^ (**Fig. 2a**). In a complementary approach, we evaluated the RPs using a quantitative total p53 assay (Alphascreen) to verify p53 expression, and observed a significant correlation between the results from both techniques (**Fig. 2b, Supplementary Fig. 3a**). In total, 77.3% of the RPs specific to the 60S and 81.3% to the 40S ribosomal subunits, when depleted, induced a ‘p53 positive’ phenotype, implying that RPs to either subunit contributed similarly to the NSP p53 response. This finding is in contrast to a study reporting that the large subunit RPs have a more profound p53 response when depleted^9^, though an arbitrary 5-fold increase in p53 was implemented as a ‘cut off’ in that study, compared to our minimum physiologically relevant 2-fold cut off which was experimentally determined. Importantly, the differential ability of the RPs when depleted to elicit p53 stabilisation was not due to the inability of the siRNA to deplete the RP mRNA and protein (**Fig. 2c, Supplementary Fig. 3b**).

**Figure 2:**
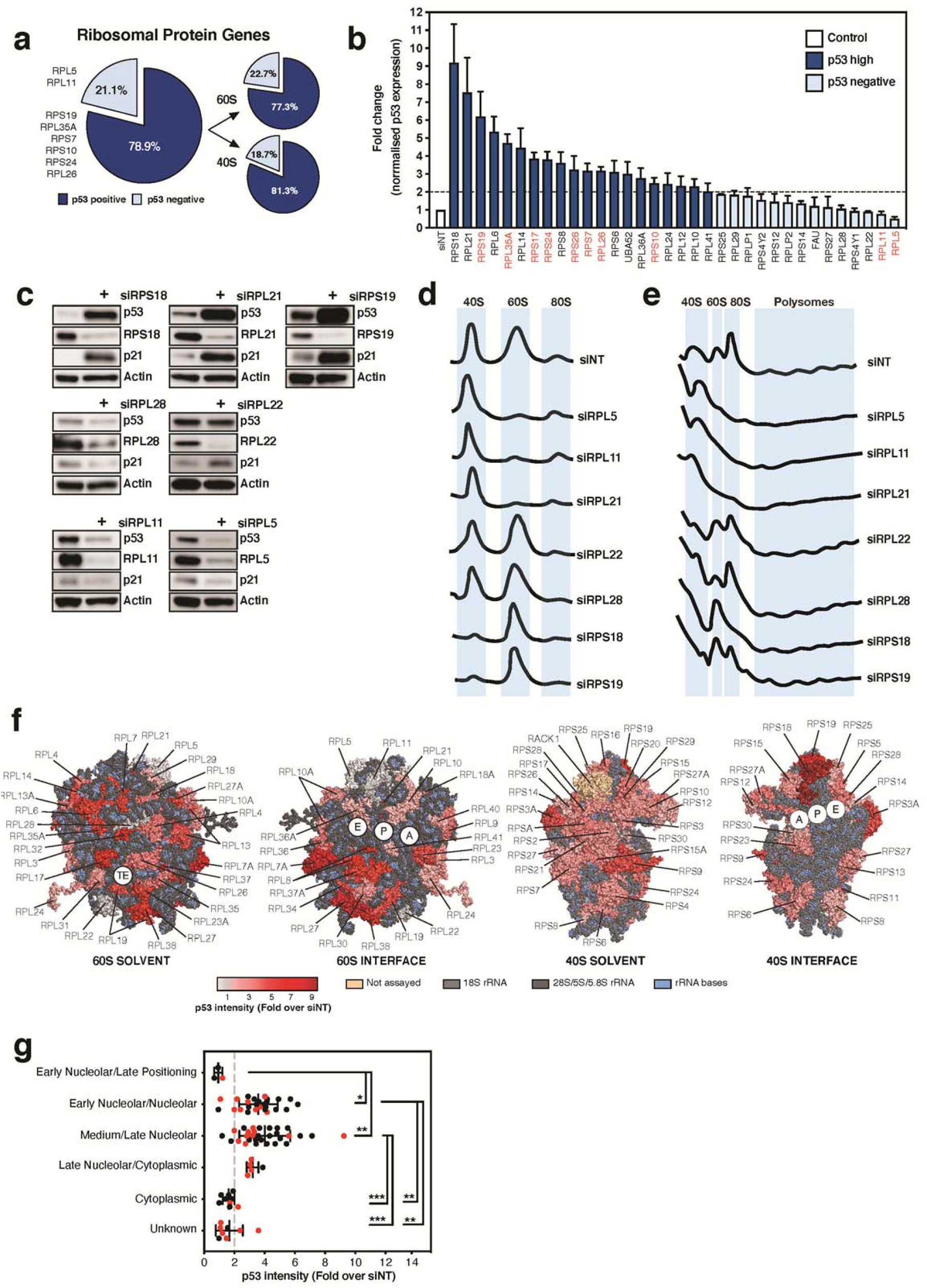
Expression of most ribosomal protein genes are integral for maintaining cellular p53 homeostasis. Given the enrichment of ribosomal protein (RP) genes in our primary screen dataset, we further investigated this group; depicted is the breakdown of screened RPs which were p53 ‘positive’ – 2-fold or greater increase in p53, and the proportion of which are located in the large (60S) or small (40S) ribosome subunit, shown in (**a**). We further verified the p53 result of approximately 50% of the RP genes (when depleted using siRNAs for 72 hours) with quantitative p53 analysis (Alphascreen) in A549 cells (note genes associated with DBA are highlighted in red) (**b**). We selected candidates which were ‘p53 positive’ (RPS18, RPL21, RPS19) and ‘p53 negative’ (RPL5, RPL11, RPL22, RPL28) to confirm knockdown at the protein level, and determined p53 and p21 protein levels using western blot analysis in A549 cells (**c**, representative blot of n=3 experiments). Cells depleted of each RP were then subjected to ribosome subunit analysis (performed under high salt conditions, **d**), to determine the effect of depletion on 60S and 40S subunits, as well as polysome analysis (**e**). We rescreened the RP genes (to incorporate those which were not assayed in the primary screen into the dataset), and mapped the p53 intensity of each RP from the screen onto the near-atomic structure of the human ribosomal 60S and 40S subunits resolved by Khatter and colleagues (PDB ID: 4UG0)^13^ (**f**), to determine if there were any patterns or regions of the ribosome where p53 intensity was focused (TE = tunnel exit, A = acceptor/aminoacyl-tRNA site, P = peptidyl-tRNA site, E = exit site). Comparison of the timing of RP incorporation into the ribosome subunit (as tabulated by de la Cruz and colleagues^14^) with p53 intensity when the RP was depleted using siRNA (**g**). Data presented as mean-/+ SD, statistical analysis: one-way ANOVA with Tukey’s multiple comparison test, \*\*\**p* < 0.001, \*\**p* < 0.01, * *p* < 0.05; red circles indicate 40S subunit RPs, black circles indicate 60S subunit RPs. Alphascreen analysis performed n=3-5 biological experiments; ribosome subunit/polysome profiles, minimum n=3 biological experiments per candidate.

We further examined whether the ability of a RP to induce the NSP correlated with the degree to which its depletion affected ribosome subunit biogenesis and function. We measured the abundance of the 40S and 60S ribosomal subunits, and the levels of mature ribosomes (80S) bound to mRNAs in polysomes following RP depletion. Consistent with the prediction, depletion of RPL21, RPS18 and RPS19, all of which induced robust stabilisation of p53, also robustly reduced the abundance of the corresponding 60S/40S subunit in which they are located, as well as the number of polysomes (**Fig. 2d & e**). In contrast, depletion of RPL22 and RPL28, which failed to induce p53 stabilisation, did not impact on 60S biogenesis, nor the number of polysomes compared to siNT (**Fig. 2d & e**). Exceptions to this were RPL5 and RPL11, whose knockdown failed to stabilise p53, even though 60S biogenesis was ablated. This observation is consistent with studies implicating ‘free’ RPL5 and RPL11 (i.e., not incorporated into a 60S) as essential for the NSP due to their ability to bind MDM2 as part of the 5S-RNP^4,10-12^.

We considered whether the location of a RP within the ribosome may predict their ability to disrupt ribosome assembly, and thus mediate p53 accumulation, upon depletion. To do this, we mapped the p53 intensity resulting from the knockdown of each RP onto the structure of the 60S and 40S subunits^13^ (**Fig. 2f**). While RPS18 and RPS19 (corresponding to two of the highest p53 intensities observed in the screen) co-located in the same region within the 40S subunit, there was no other clear evidence supporting that the specific location of a RP in the ribosome would increase p53 stabilisation if depleted. Finally, we tested the hypothesis that RPs which integrate early into their respective ribosomal subunit (i.e., within the nucleolus) might be essential for the core structure, thus when depleted, would have the most profound effect on ribosome assembly and the NSP. By comparing p53 intensity and the published timing of integration of each RP into the ribosome^14^ (**Supplementary File 5**), we demonstrated that the p53 levels were significantly higher following knockdown of those RPs which integrate into their respective subunits during early nucleolar stages of ribosome assembly (**Fig. 2g**). Thus, the ability of RPs to stabilise p53 correlated with their propensity to cause significant disruption to ribosome subunit assembly when depleted. This may explain, at least in part, why not all components of the ribosome, when mutated or deleted, contribute to ribosomopathies. For example, RPL22 and RPL28, which do not perturb ribosome subunit assembly when depleted, have not been associated with DBA to date.

Having identified the major classes of genes, including RPs, whose deletion leads to stabilisation of p53, we next determined the role of the NSP in this process; *a priori*, we predicted that only those genes directly involved in RiBi would be dependent on the NSP to stabilise p53 when depleted. To address this question, we took an unbiased approach to identify the key components of NSP that can be targeted to inactivate NSP-mediated p53 stabilisation. Accordingly, we performed a genome-wide RNAi screen to determine the genes whose depletion suppressed p53 accumulation in response to nucleolar stress induced by knockdown of RPS19, the prototypical DBA gene known to induce NSP when depleted ^15-17^ (termed ‘modifiers of ribosomal stress’ screen; **Fig. 3a & Supplementary File 6**). Using a cut-off for normalised p53 intensity of up to and including 0.5 (log_2_=-1, calculated based on 3 standard deviations (SD) above the positive control, siTP53+siRPS19), we identified 64 genes essential for a functional NSP (**Fig. 3a & Supplementary Fig. 4a**). We rescreened these 64 candidates (outlined in Methods), to identify candidates which recapitulated the primary screen phenotype (i.e. suppressed p53 response when co-depleted with siRPS19) with two or more siRNA duplexes. Critically, in addition to TP53, both RPL5 and RPL11 were the top ranked candidates which, when depleted, reduced p53 accumulation in response to NSP activation, while no other RPs reached this cut-off. This observation is in contrast to previous reports suggesting a variety of RPs regulate p53 stability (e.g. RPL23^18,19^, RPL26^20,21^, RPS3^22^, RPS7^23^, RPS14^24^, RPS25^25^, RPS27A^26^, RPS27 and RPS27L^27^, RPS15, RPS20 and RPL37^28^). Our study, therefore, functionally defines RPL5 and RPL11 as the only RPs essential for the NSP, consistent with their proposed role in the 5S-RNP interaction with MDM2. Similarly, non-RP factors previously reported to be linked to RiBi and p53 activity (e.g. SRSF1, GLTSCR2 (PICT1), HEXIM1, MYBBP1A, RRP8 (NML) and NPM1^29-34^) were not identified as high-ranking candidates, suggesting, at least in this system under the kinetics used for the assays, they are not essential for NSP-induced stabilisation of p53 and/or may play tissue or developmentally-specific roles in the NSP.

**Figure 3:**
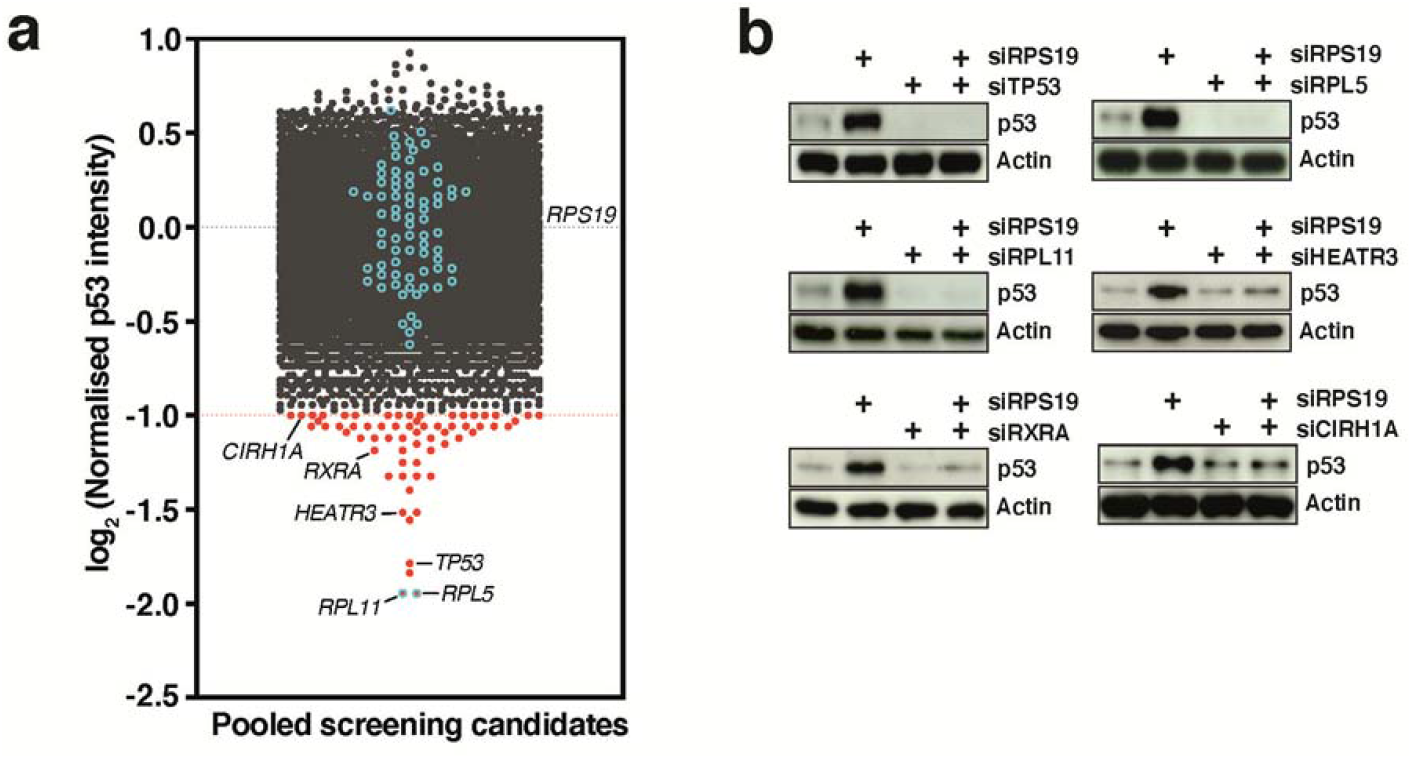
High-throughput screening for modifiers of ribosomal stress due to activation of the canonical nucleolar surveillance pathway (NSP). In a similar approach (outlined in Fig. 1A), we performed a genome-wide RNAi screen to identify modifiers of ribosomal stress, by co-depleting RPS19 with every gene in the genome. After conducting the screen, candidates were further triaged using gene-expression data from RNAseq analysis of A549 cells depleted of RPS19 (RPKM cutoff of 0.05 or greater) to yield 14,577 ‘expressed’ screen candidates. The ‘expressed’ screen candidates were then graphed normalised to RPS19 depletion (**a**); candidates in red are those with a log_2_ value of ≤-1 (total 64 candidates). Ribosomal protein (RP) genes in the screening data are demarcated with blue circles. A selection of these candidates (TP53, RPL5, RPL11, HEATR3, RXRA and CIRH1A) were then further subjected to candidate-based validation in A549 cells (**b**); by co-depletion of candidates with siRPS19 for 72 hours and analysed by western blotting (representative of n=3 biological experiments).

In addition to TP53, RPL5 and RPL11, we further validated a selection of candidates from the screen including HEATR3, RXRA and CIRH1A as *bone fide* modulators of the p53 response (**Fig. 3b, Supplementary Fig. 4a, c-h & i-n**). HEATR3 (HEAT-repeat containing 3) was of significant interest as a novel direct regulator of the NSP and the 5S-RNP-MDM2 axis, as bioinformatic domain alignment suggested that HEATR3 is a human homolog of yeast symportin 1 (Syo1) protein, which enables import of rpL5 and rpL11 into the nucleus of *Saccharomyces cerevisiae*^35,36^, and acts as a scaffold for 5S-RNP biogenesis prior to incorporation into the pre-60S ribosomal subunit^35^. To analyse any structural similarities between the human and yeast proteins, we modelled the HEATR3 structure based on the *Chaetomium thermophilium* Syo1 (ctSyo1) crystal structure^36^ using ‘Modeller’ (**Fig. 4a & Supplementary Fig. 5**), which indicates the potential for RPL5 and RPL11 binding on opposite sides of the HEAT repeats similar to that shown for ctSyo1^35,36^. In support of this model, co-immunoprecipitation experiments from A549 cells co-transfected with myc-tagged HEATR3 (MT-HEATR3) and either FLAG-tagged RPL5 or RPL11 confirmed that RPL11 and RPL5 bind to HEATR3 *in situ* (**Fig. 4b**). Moreover, depletion of HEATR3 partially phenocopied RPL5 and RPL11 knockdown, resulting in a marked reduction in 60S subunit production (**Fig. 4c**), the number of polysomes (**Fig. 4d**) and 5S-RNP binding to MDM2 (**Fig. 4e & f**). The reduced efficacy of HEATR3 depletion to disrupt RiBi and NSP compared to RPL5 and RPL11 suggests there may also be HEATR3-independent pathways by which RPL5 and RPL11 can assemble into 5S-RNP. Even so, *in toto*, these findings strongly suggest that HEATR3 is a functional homolog of Syo1 and important for 60S assembly and NSP in human cells through its ability to interact with the 5S-RNP (**Fig. 4g**).

**Figure 4:**
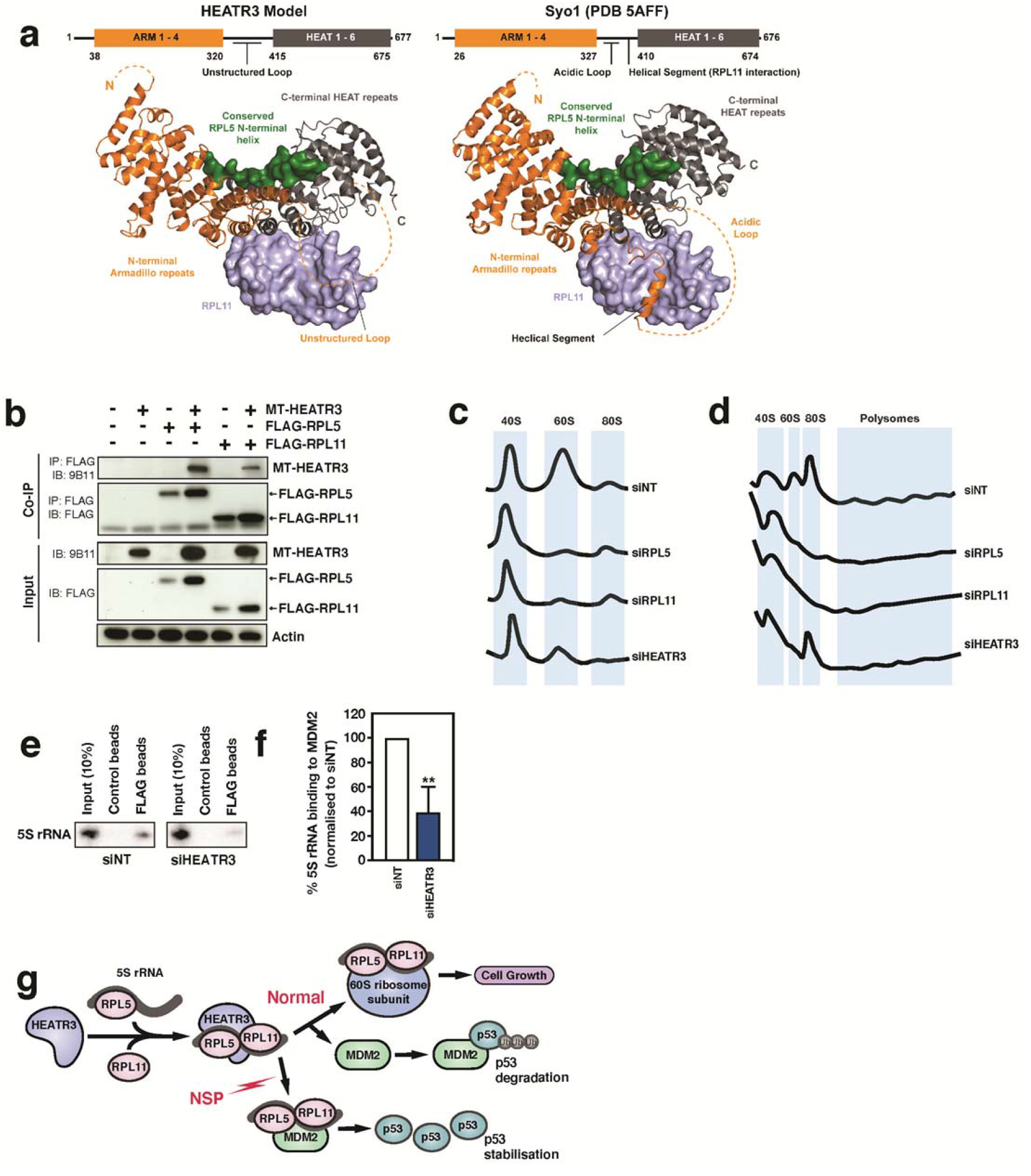
Analysis of the HEAT-repeat containing 3 (HEATR3) protein and its role in ribosome and 5S-RNP biogenesis. Comparison between *C. thermophilium* Syo1 (ctSyo1) (PDB 5AFF)^36^ and the predicted human HEATR3 structure (**a**). HEATR3 secondary structure and domain modelling indicates the presence of an N-terminal Armadillo (ARM, orange), and a C-terminal HEAT repeat domain (dark grey), similar to the yeast Symportin 1 (Syo1) protein. A domain schematic (to scale; top) and a cartoon model (bottom) are shown for each protein. Similar to the ctSyo1 protein, HEATR3 contains four N-terminal Armadillo (ARM) repeats and six C-terminal HEAT repeats. In the case for HEATR3, these regions are connected by a central, long and unstructured loop, whereas ctSyo1 has an acidic loop with a helical segment (Glu389 to Gly399) likely responsible, at least in part, for the binding of rpL11 (light blue, surface representation) to the protein^35^. A conserved N-terminal segment of RPL5 (green, surface representation) may also interact with HEATR3 (similar to Syo1). Co-immunoprecipitation (CoIP) analysis of human myc-tagged HEATR3 (MT-HEATR3) with FLAG-tagged human RPL5 and RPL11 proteins in HEK293 cells (**b**). Ribosome (**c**) and polysome (**d**) profiling analysis of A549 cells depleted of RPL5, RPL11 or HEATR3 (and non-targeting siRNA, siNT) for 72 hours (note that the NT, RPL5 and RPL11 data traces presented here are already presented in Fig 2d and e, and are replicated in this figure to directly compare the effect of HEATR3 depletion with these conditions). Northern blot analysis of the association of MDM2 with 5S rRNA after 48-hour HEATR3 depletion in U2OS cells expressing FLAG-MDM2 (**e**) and quantitation (**f**). Schematic of the predicted role of HEATR3 in 5S-RNP biogenesis (“Normal”) and the NSP (**g**). Error bars represent SD and statistical analysis performed using unpaired student t-test (** *p* < 0.01, n=3 experiments).

Having functionally defined RPL5, RPL11 and HEATR3 as direct regulators of the NSP, we next used their depletion (RNAi) to determine how important a functional NSP is for stabilisation of p53 by stresses not traditionally implicated in RiBi. To do this, a representative selection of the 827 genes identified as ‘p53 positive’ (i.e., whose depletion increased p53 levels; **Fig. 1b&c**; 232 genes representing nucleolar, ribosome, splicing, Pol II, proteasome, cell cycle and other gene classes) were rescreened to determine if their ability to stabilise p53 when depleted was dependent on the NSP (**Fig. 5 & Supplementary File 7)**. As expected, the ability of RP and other nucleolar/RiBi-related genes to robustly activate p53 when depleted was blocked when the NSP was inactivated by co-depletion of either RPL5, RPL11 or HEATR3. Notably, the overall effect of HEATR3 depletion to reduce p53 accumulation was less profound than RPL5 and RPL11 depletion, and for a subset of large RNPs, HEATR3 depletion completely failed to block induction of p53 (**Fig. 5a**&**b, Supplementary File 7**). Thus, HEATR3 is necessary for 5S-RNP-MDM2 complex assembly in response to the disruption of many (but not all) RiBi proteins, consistent with the observations above that HEATR3-independent pathways by which RPL5 and RPL11 can assemble into 5S-RNP may exist.

**Figure 5:**
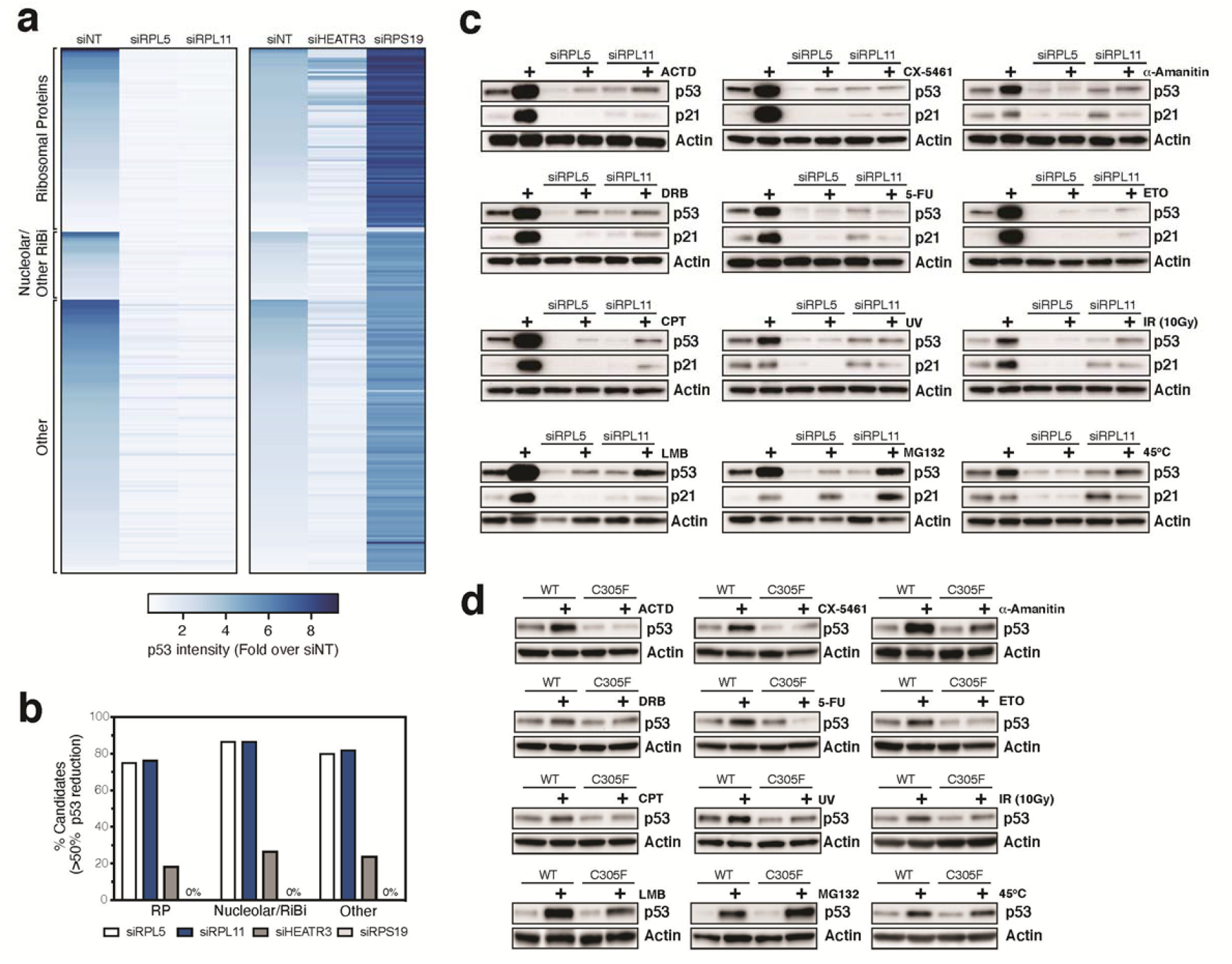
The NSP, via RPL5 and RPL11, is required to stabilise p53 in response to broad range of genetic, pharmacological and pathophysiologic stresses. To identify which candidate genes are required for the canonical NSP, we rescreened a selection of candidates from the primary p53 stabilisation screen described in Fig. 3 (ribosomal proteins, nucleolar/RiBi candidates and “other” p53 positive candidates, 232 genes in total) in the presence of non-targeting, RPL5, RPL11, HEATR3 or RPS19 siRNAs in A549 cells (**a**). From this analysis, we further quantified the number of candidates screened from each group (ribosomal proteins, nucleolar/RiBi and other) for which their p53 response could be suppressed by ≥ 50% when co-depleted with RPL5, RPL11, HEATR3 or RPS19 siRNAs (**b**). We further tested a panel of pharmacological agents and pathophysiological stressors when A549 cells were depleted of RPL5 and RPL11 for 48 hours. Cells were treated for 24 hours with pharmacological agents Actinomycin D (ACTD, 5 nM), α-Amanitin (2.5 μM), Doxorubicin (DRB, 500 nM), 5-Fluoruracil (5-FU, 50 μM), Etoposide (ETO, 50 μM), Camptothecin (CPT, 50 nM), Leptomycin B (LMB, 10 ng/mL) or MG132 (10 μM). Alternatively, cells were treated with pathophysiological stressors UV (50 J/m^2^), gamma irradiation (10Gy), or subjected to heat shock (45°C, 30 minutes), then incubated at 37°C for 3 hours post treatment. At the end of treatment, protein was harvested and subjected to western blot analysis for p53 and p21 protein expression (**c**). The same panel of stressors (and treatment conditions) were testing on mouse embryonic fibroblasts (MEFs) isolated from either Mdm2 wild-type (WT) or mice homozygous for the Mdm2 C305F mutation (C305F) to determine p53 expression (**d**). n=3-4 biological replicates for each condition, the most representative experiment for each treatment is presented.

Critically, and unexpectedly, the ability of major classes of genes not traditionally associated with the ribosome or the nucleolus (e.g., RNA splicing, cell cycle and Pol II, **Fig.1c**) to stabilise p53 following their depletion was also ablated upon co-knockdown of RPL5 or RPL11, and to a lesser degree HEATR3 (**Fig. 5a & b, Supplementary File 7**). The p53 suppression was not simply due to reduced ribosome assembly (and therefore reduced p53 mRNA translation) as a consequence of RPL5 or RPL11 depletion (**Fig. 4c & d**), because co-depletion of RPS19 failed to blunt p53 accumulation, despite RPS19 depletion causing a similar defect in ribosomal subunit assembly and polysomes (**Fig. 2d & e**). Together, these data suggest the NSP is required for robust stabilisation of p53 in response to the dysregulation of a large number of eukaryotic genes and cellular processes that are not traditionally associated with RiBi.

Given these unexpected findings, we extended these studies to determine the requirement of a functional NSP to mediate stabilisation of p53 in response to a broad range of pharmacological agents and pathophysiologic stressors, including inhibitors of Pol I & II, nucleic acid synthesis inhibitors, agents that induce DNA damage, nuclear export inhibitors and proteotoxic stress.cIntriguingly, inactivation of the NSP by either RPL5 or RPL11 depletion ablated the ability of Pol I/II inhibitors, nuclei acid synthesis inhibitors and all classes of DNA damage-inducing agents to stabilise p53. In contrast, the ability of proteotoxic stresses including proteasomal inhibitors, nuclear transport inhibitors and heat shock to increase p53 levels were only moderately, or not at all blunted by inactivation of the NSP (**Fig. 5c & Supplementary Fig. 6b**). We also confirmed these findings using a high-content screening-based approach (**Supplementary Fig. 6c**), where HEATR3 depletion also blunted the response, however, not as efficiently as RPL5/L11 depletion. We noted that knockdown of RPL5 was consistently more potent at blocking the NSP compared to RPL11 or HEATR3, suggesting RPL5 may modulate p53 by mechanisms in addition to inhibitory binding of 5S-RNP to MDM2. Consistent with this, we observed that knockdown of RPL5 but not RPL11 significantly reduced p53 mRNA levels (**Supplementary Fig. 6d**), although the mechanism of this reduction was not investigated further.

To further validate our results in a model of NSP inactivation (other than RPL5 and RPL11 depletion), we used embryonic fibroblasts (MEFs) isolated from mice harbouring the Mdm2^C305F^ mutation, which disrupts RPL5 and RPL11 (ergo 5S-RNP) binding to MDM2, thereby inactivating the NSP^4,12^. We again tested a complement of nuclear and physiological stressors (**Fig. 5d & Supplementary Fig. 6e**) in these cells. Consistent with RPL5/RPL11 knockdown, the Mdm2^C305F^ mutation prevented the p53 response upon exposure to Pol I/II inhibitors, nucleic acid synthesis inhibitors and all classes of DNA damage inducing agents, but not proteotoxic stress. Taken together, the data indicates that an intact NSP is required for the stabilisation of p53 in response to a broad range of cellular stresses, not just ribosomal/nucleolar stress. Notably, the quantitative effect of Mdm2^C305F^ mutation to blunt p53 accumulation in response to stress more closely reflected the effect of RPL11 depletion than RPL5 depletion, consistent with the conclusions above that RPL5 may modulate p53 by mechanisms in addition to inhibitory binding of 5S-RNP to MDM2.

In summary, using global screening approaches, we have identified the complement of genes and pathways functionally required for stabilisation of p53 in response to the canonical NSP. Our data definitively demonstrate that RPL5 and RPL11 do not induce p53 stabilisation when depleted, and are the only RPs essential for functional NSP to stabilise p53^4,12,37^. Furthermore, we demonstrate that one of the top hits, HEATR3, is a potential ribosome assembly factor required for mammalian 60S ribosomal subunit assembly through binding of RPL5 and RPL11. Consistent with an essential role for HEATR3 in NSP-mediated stabilisation of p53, HEATR3 depletion leads to reduced association of the 5S-RNP with MDM2.

Critically, by inactivating the NSP, we demonstrate that pharmacological agents and pathophysiological conditions leading to genotoxic stress, as well as the majority of genes whose loss-of-function stabilises p53, do so in an NSP-dependent fashion. Our data provides experimental support to Rubbi and Milner’s original hypothesis that the nucleolus, through the NSP, is a universal stress sensor responsible for p53 homeostasis within cells^38^. Thus, we conclude that the well-described mechanisms of genotoxic stress which induce extensive post-translational regulation of p53, thereby modulating its interaction with MDM2, are insufficient in the absence of a functioning NSP to robustly stabilise p53. The exception to this rule appears to be pathological conditions and pharmacologic agents that result in proteasomal stress, which stabilise p53 largely independently of the NSP. This is consistent with MDM2-mediated degradation of p53 being dependent on a functioning proteasome to degrade ubiquitinated p53.

The differential ability of ribosome components to induce p53 stabilisation following their depletion correlated directly with the degree of disruption of RiBi and ribosome assembly. By extrapolation, we propose that all nuclear acting-pathological conditions,-pharmacologic agents or genetic inactivating lesions stabilise p53 in a 5S-RNP-MDM2 dependent fashion, through disruption of RiBi. Consistent with this, ribosomal DNA (rDNA) is highly sensitive to DNA damage (a single lesion in the rDNA is sufficient to cause cell cycle arrest^39^), and most cytotoxic drugs and pathologic conditions that induce DNA damage have been reported to cause defects in RiBi. We propose that the nucleolus functions as the cellular equivalent of a sentinel or “canary in the coal mine” to detect a broad range of cellular stresses and mediate stabilisation of p53. In this model, the nucleolus acts a sensitivity gauge, whereby stresses such as DNA damage can only stabilise p53 if the stress is of sufficient magnitude to perturb RiBi/nucleolar function, thereby preventing minor cellular insults from inappropriately inhibiting proliferation. Given that RiBi is the most energy-expensive process a cell undertakes, the evolution of such a mechanism also ensures RiBi remains hardwired to proliferative capacity through p53 activity.

Finally, due to the central role RiBi and the NSP plays in the regulation of p53, we suggest a paradigm-shift in thinking is required for how this axis contributes to cancer pathogenesis. Due to the pervasive stress tumour cells are exposed to, we propose that overcoming NSP-induced p53 activation is likely to be a very frequent step in malignant transformation. Indeed, RP genes are hemizygously deleted in 43% of human cancers, and almost always in concert with *TP53* mutations, while such RP deletions are infrequent in *TP5*3-intact tumours^40,41^. This is consistent with chronic activation of the NSP in response to RP deletion being incompatible with malignant transformation and negatively selected for unless p53 is inactivated.

## Supporting information

Supplementary Data File 1 - p53 stabilisation screen

Supplementary Data File 2 - STRING analysis

Supplementary Data File 3 - Gene Ontology Analysis

Supplementary Data File 4 - LOCATE sub cellular localisation analysis

Supplementary Data File 5 - assembly analysis

Supplementary Data File 6 - modifiers of ribosomal stress screen

Supplementary Data File 7 - heatmap data (corresponding to Figure 5)

Supplementary Data File 8 - high content screening image analysis pipelines

Supplementary Information (including Methods and Supplementary Figures)

## Acknowledgements

Thanks to the Captain Courageous Foundation (captaincourageous.com.au) in particular, Jessica and Jeff Bond, and Bill Steele for their ongoing support and funding for this project. We would like to acknowledge the founding and current members of the Australian Diamond Blackfan Anaemia (ADBA) program, consisting of Sheren J. Al-Obaidi, Luen Bik To, Sarah C.E. Bray, Richard J. D’Andrea, Jianmin Ding, Amee J. George, Thomas J. Gonda, Ross D. Hannan, Melissa Ilsley, S. Peter Klinken, Lorena Núñez-Villacís, Richard B. Pearson, Ben Saxon, Hamish S. Scott, Adam Stephenson, Adam Stevenson, Parvathy Venugopal, Amilia Wee, Louise N. Winteringham and Mei Szin Wong. Our sincerest thanks go to Yanping Zhang (UNC Chapel Hill, USA) for providing the Mdm2 mouse strain used in this work. We would also like to thank George Thomas (IDIBELL, Spain) for his valuable discussions.

We would like to thank the Imaging and Cytometry Facility (Dr Harpreet Vohra and Mr Michael Devoy), the ANU Bioinformatics Consultancy (Dr Zhi-Ping Feng and Mr Cameron Jack), and the staff at the Australian Phenomics Facility, the ACRF Biomolecular Resource Facility and the ANU Centre for Therapeutic Discovery at the John Curtin School of Medical Research at the Australian National University for their assistance. We would also like to thank the ACRF Victorian Centre for Functional Genomics (Ms Jennii Luu and Mr Daniel Thomas), the Molecular Genomics Core (Dr Gisela Mir Arneau and Ms Aga Borcz), the Bioinformatics Core (Mr Jason Ellul) and the Advanced Centre for Cancer Cell Isolation and Flow Cytometry (Ms Viki Milovac and Ms Sophie Curcio) located at the Peter MacCallum Cancer Centre for their technical assistance, and all members of the Engel lab, especially Philipp Becker and Mona Höcherl.

This work was supported by funding from the Captain Courageous Foundation and the National Health and Medical Research Council (NHMRC) of Australia (Project Grants #1100654, #1158732, #1102609, Program Grant #1053792 and Senior Research Fellowships to R.D.H (#1116999) and R.B.P (#1058586). C.E. is supported by the ‘Emmy-Noether-Programme’, DFG grant no. EN 1204/1-1. U.K. was supported by the Swiss National Science Foundation (SNSF) (grant 31003A_166565 and the NCCR ‘RNA and disease’). N.J.W is supported by funding from the DBAF, DBA UK and the BBSRC (BB/R00143X/1). A.J.G is supported by a Captain Courageous Foundation Fellowship.

The Victorian Centre for Functional Genomics (K.J.S.) is funded by the Australian Cancer Research Foundation (ACRF) and Phenomics Australia, through funding from the Australian Government’s National Collaborative Research Infrastructure Strategy (NCRIS) program and the Peter MacCallum Cancer Centre Foundation. The ANU Centre for Therapeutic Discovery (A.J.G) is funded by the ACRF, Australian National University and ACT Health.

## Author Contributions

K.M.H, R.D.H and A.J.G conceived and designed the study and wrote the manuscript; all other authors have reviewed and edited the manuscript. K.J.S, R.D.H and A.J.G. designed the high-throughput screening approaches utilised in this study. A.J.G, P.S, J.H and K.M.P. performed the high-throughput screens/high-content imaging assays and developed the image analysis pipelines. M.E., L.K.S, Z-P.F, C.M.G, P.B.M, K.J.S and A.J.G performed the analysis and visualisation of screening data. M.S.W. and A.J.G designed and executed the validation of screening candidates and HEATR3 coimmunoprecipitation analysis. J.K.L and N.J.W. performed the co-immunoprecipitation analysis of 5S rRNA with MDM2. C.M., U.K and A.J.G performed the ribosome shading and assembly analyses. C.E. performed the HEATR3/Syo1 sequence alignments and predictive modelling. K.M.H., P.S, M.S.W, T.D.H, and P.P. performed the ribosomal subunit and polysomal analyses. P.S, M.S.W, N.H, K.D.W, T.D.W, S.J.A, L.N.V, P.P, M.P, G.B, R.F and A.J.G performed the molecular biology and biochemical experiments. J.F, T.J.G., K.J.S, U.K., R.B.P., N.J.W., R.D.H. and A.J.G. contributed valuable interpretation and academic discussion. R.D.H. and A.J.G. co-supervised the project.

## References

1 Zilfou, J. T. & Lowe, S. W. Tumor suppressive functions of p53. Cold Spring Harb Perspect Biol 1, a001883, doi:10.1101/cshperspect.a001883 (2009).

2 Dai, M. S. et al.. Physical and functional interaction between ribosomal protein L11 and the tumor suppressor ARF. J Biol Chem 287, 17120–17129, doi:10.1074/jbc.M111.311902 (2012).

3 Boulon, S., Westman, B. J., Hutten, S., Boisvert, F. M. & Lamond, A. I. The nucleolus under stress. Mol Cell 40, 216–227, doi:10.1016/j.molcel.2010.09.024 (2010).

4 Sloan, K. E., Bohnsack, M. T. & Watkins, N. J. The 5S RNP couples p53 homeostasis to ribosome biogenesis and nucleolar stress. Cell Rep 5, 237–247, doi:10.1016/j.celrep.2013.08.049 (2013).

5 Donati, G., Peddigari, S., Mercer, C. A. & Thomas, G. 5S ribosomal RNA is an essential component of a nascent ribosomal precursor complex that regulates the Hdm2-p53 checkpoint. Cell Rep 4, 87–98, doi:10.1016/j.celrep.2013.05.045 (2013).

6 von Mering, C. et al.. STRING: a database of predicted functional associations between proteins. Nucleic Acids Res 31, 258–261, doi:10.1093/nar/gkg034 (2003).

7 Sprenger, J. et al.. LOCATE: a mammalian protein subcellular localization database. Nucleic Acids Res 36, D230–233, doi:10.1093/nar/gkm950 (2008).

8 Ulirsch, J. C. et al.. The Genetic Landscape of Diamond-Blackfan Anemia. Am J Hum Genet 103, 930–947, doi:10.1016/j.ajhg.2018.10.027 (2018).

9 Nicolas, E. et al.. Involvement of human ribosomal proteins in nucleolar structure and p53- dependent nucleolar stress. Nat Commun 7, 11390, doi:10.1038/ncomms11390 (2016).

10 Bursac, S. et al.. Mutual protection of ribosomal proteins L5 and L11 from degradation is essential for p53 activation upon ribosomal biogenesis stress. Proc Natl Acad Sci U S A 109, 20467–20472, doi:10.1073/pnas.1218535109 (2012).

11 Fumagalli, S., Ivanenkov, V. V., Teng, T. & Thomas, G. Suprainduction of p53 by disruption of 40S and 60S ribosome biogenesis leads to the activation of a novel G2/M checkpoint. Genes Dev 26, 1028–1040, doi:10.1101/gad.189951.112 (2012).

12 Macias, E. et al.. An ARF-independent c-MYC-activated tumor suppression pathway mediated by ribosomal protein-Mdm2 Interaction. Cancer Cell 18, 231–243, doi:10.1016/j.ccr.2010.08.007 (2010).

13 Khatter, H., Myasnikov, A. G., Natchiar, S. K. & Klaholz, B. P. Structure of the human 80S ribosome. Nature 520, 640–645, doi:10.1038/nature14427 (2015).

14 de la Cruz, J., Karbstein, K. & Woolford, J. L., Jr. Functions of ribosomal proteins in assembly of eukaryotic ribosomes in vivo. Annu Rev Biochem 84, 93–129, doi:10.1146/annurev-biochem-060614-033917 (2015).

15 Jaako, P. et al.. Disruption of the 5S RNP-Mdm2 interaction significantly improves the erythroid defect in a mouse model for Diamond-Blackfan anemia. Leukemia 29, 2221–2229, doi:10.1038/leu.2015.128 (2015).

16 Jaako, P. et al.. Mice with ribosomal protein S19 deficiency develop bone marrow failure and symptoms like patients with Diamond-Blackfan anemia. Blood 118, 6087–6096, doi:10.1182/blood-2011-08-371963 (2011).

17 Sjogren, S. E. et al.. Glucocorticoids improve erythroid progenitor maintenance and dampen Trp53 response in a mouse model of Diamond-Blackfan anaemia. Br J Haematol 171, 517–529, doi:10.1111/bjh.13632 (2015).

18 Dai, M. S. et al.. Ribosomal protein L23 activates p53 by inhibiting MDM2 function in response to ribosomal perturbation but not to translation inhibition. Mol Cell Biol 24, 7654–7668, doi:10.1128/MCB.24.17.7654-7668.2004 (2004).

19 Jin, A., Itahana, K., O’Keefe, K. & Zhang, Y. Inhibition of HDM2 and activation of p53 by ribosomal protein L23. Mol Cell Biol 24, 7669–7680, doi:10.1128/MCB.24.17.7669-7680.2004 (2004).

20 Ofir-Rosenfeld, Y., Boggs, K., Michael, D., Kastan, M. B. & Oren, M. Mdm2 regulates p53 mRNA translation through inhibitory interactions with ribosomal protein L26. Mol Cell 32, 180–189, doi:10.1016/j.molcel.2008.08.031 (2008).

21 Zhang, Y. et al.. Negative regulation of HDM2 to attenuate p53 degradation by ribosomal protein L26. Nucleic Acids Res 38, 6544–6554, doi:10.1093/nar/gkq536 (2010).

22 Yadavilli, S. et al.. Ribosomal protein S3: A multi-functional protein that interacts with both p53 and MDM2 through its KH domain. DNA Repair (Amst) 8, 1215–1224, doi:10.1016/j.dnarep.2009.07.003 (2009).

23 Chen, D. et al.. Ribosomal protein S7 as a novel modulator of p53-MDM2 interaction: binding to MDM2, stabilization of p53 protein, and activation of p53 function. Oncogene 26, 5029–5037, doi:10.1038/sj.onc.1210327 (2007).

24 Zhou, X., Hao, Q., Liao, J., Zhang, Q. & Lu, H. Ribosomal protein S14 unties the MDM2- p53 loop upon ribosomal stress. Oncogene 32, 388–396, doi:10.1038/onc.2012.63 (2013).

25 Zhang, X. et al.. Identification of ribosomal protein S25 (RPS25)-MDM2-p53 regulatory feedback loop. Oncogene 32, 2782–2791, doi:10.1038/onc.2012.289 (2013).

26 Sun, X. X., DeVine, T., Challagundla, K. B. & Dai, M. S. Interplay between ribosomal protein S27a and MDM2 protein in p53 activation in response to ribosomal stress. J Biol Chem 286, 22730–22741, doi:10.1074/jbc.M111.223651 (2011).

27 Xiong, X., Zhao, Y., He, H. & Sun, Y. Ribosomal protein S27-like and S27 interplay with p53-MDM2 axis as a target, a substrate and a regulator. Oncogene 30, 1798–1811, doi:10.1038/onc.2010.569 (2011).

28 Daftuar, L., Zhu, Y., Jacq, X. & Prives, C. Ribosomal proteins RPL37, RPS15 and RPS20 regulate the Mdm2-p53-MdmX network. PLoS One 8, e68667, doi:10.1371/journal.pone.0068667 (2013).

29 Fregoso, O. I., Das, S., Akerman, M. & Krainer, A. R. Splicing-factor oncoprotein SRSF1 stabilizes p53 via RPL5 and induces cellular senescence. Mol Cell 50, 56–66, doi:10.1016/j.molcel.2013.02.001 (2013).

30 Kumazawa, T. et al.. Novel nucleolar pathway connecting intracellular energy status with p53 activation. J Biol Chem 286, 20861–20869, doi:10.1074/jbc.M110.209916 (2011).

31 Kuroda, T. et al.. RNA content in the nucleolus alters p53 acetylation via MYBBP1A. EMBO J 30, 1054–1066, doi:10.1038/emboj.2011.23 (2011).

32 Lew, Q. J. et al.. Identification of HEXIM1 as a positive regulator of p53. J Biol Chem 287, 36443–36454, doi:10.1074/jbc.M112.374157 (2012).

33 Sasaki, M. et al.. Regulation of the MDM2-P53 pathway and tumor growth by PICT1 via nucleolar RPL11. Nat Med 17, 944–951, doi:10.1038/nm.2392 (2011).

34 Lindstrom, M. S. NPM1/B23: A Multifunctional Chaperone in Ribosome Biogenesis and Chromatin Remodeling. Biochem Res Int 2011, 195209, doi:10.1155/2011/195209 (2011).

35 Calvino, F. R. et al.. Symportin 1 chaperones 5S RNP assembly during ribosome biogenesis by occupying an essential rRNA-binding site. Nat Commun 6, 6510, doi:10.1038/ncomms7510 (2015).

36 Kressler, D. et al.. Synchronizing nuclear import of ribosomal proteins with ribosome assembly. Science 338, 666–671, doi:10.1126/science.1226960 (2012).

37 Teng, T., Mercer, C. A., Hexley, P., Thomas, G. & Fumagalli, S. Loss of tumor suppressor RPL5/RPL11 does not induce cell cycle arrest but impedes proliferation due to reduced ribosome content and translation capacity. Mol Cell Biol 33, 4660–4671, doi:10.1128/MCB.01174-13 (2013).

38 Rubbi, C. P. & Milner, J. Disruption of the nucleolus mediates stabilization of p53 in response to DNA damage and other stresses. EMBO J 22, 6068–6077, doi:10.1093/emboj/cdg579 (2003).

39 van Sluis, M. & McStay, B. A localized nucleolar DNA damage response facilitates recruitment of the homology-directed repair machinery independent of cell cycle stage. Genes Dev 29, 1151–1163, doi:10.1101/gad.260703.115 (2015).

40 Ajore, R. et al.. Deletion of ribosomal protein genes is a common vulnerability in human cancer, especially in concert with TP53 mutations. EMBO Mol Med 9, 498–507, doi:10.15252/emmm.201606660 (2017).

41 Fancello, L., Kampen, K. R., Hofman, I. J., Verbeeck, J. & De Keersmaecker, K. The ribosomal protein gene RPL5 is a haploinsufficient tumor suppressor in multiple cancer types. Oncotarget 8, 14462–14478, doi:10.18632/oncotarget.14895 (2017).

