## Supplementary Data File 8 - high content screening image analysis pipelines for "Nuclear stabilisation of p53 requires a functional nucleolar surveillance pathway"

### p53_TA.V3_20x_SCREEN (Cellomics ArrayScan VTi)

Last Modified on 8/30/2013 11:53:28 AM

*NOTE: The information here represents data that is saved in the database with an Assay Protocol. This data does not include scan settings like Field Offset, Form Factor, Scan Area, Store Images or Show Composite Image.*

**Comments**

Event1=(None) Event2=(None) Event3=(None)

| Image Acquisition | |
|---|---|
| Objective | 20x |
| Camera Name | ORCA-ER;1.00 |
| Acquisition Camera Mode | Standard (1024x1024;2x2) |
| AutoFocus Camera Mode | AutoFocus (1024x1024;4x4) |
| AutoFocus Field Interval | 2 |

| AutoFocus Parameters | |
|---|---|
| Fine Focus Step Size | 9.9 |
| Fine Focus Plane Count | 9 |
| Coarse Focus Step Size | 39.6 |
| Coarse Focus Plane Count | 16 |
| Smart Focus Plane Count | 21 |
| Use Extended Range Focusing | False |
| Apply Backlash Correction | False |
| AutoFocus Method | STANDARD |
| Use Relaxed Pass/Fail Criteria | False |
| Focus Edge Threshhold | 0 |
| Focus Adjustment | 0 |
| Focus Score Min Ratio | 0.25 |
| Focus Score Mid Ratio | 0.4 |
| Focus Score Max Ratio | 0.5 |
| Focus Exposure Time for AutoExpose (seconds) | 0.02 |

| Scan Limits | |
|---|---|
| Max Fields for Well | 25 |
| Min Objects for Well | No Limit |
| Max Sparse Fields for Well | No Limit |
| Min Objects for Field | N/A |
| Max Sparse Wells for Plate | N/A |

| Channel 1: DAPI | | |
|---|---|---|
| Dye | XF93 - Hoechst | |
| Apply Illumination Correction | False | |
| Apply Background Correction | True | |
| Gain | 100 | |
| Z Offset | 0.00 | |
| Step Size | 0.00 | |
| Number of Steps | 0 | |
| Projection Method | | |
| Projection Direction | None | |
| Detection Mode | Widefield | |
| Grid Type | | |
| Pin Hole Size | | |
| Intensity Percent | 100 | |
| **Exposure Parameters** | | |
| Method | Fixed | |
| Exposure Time (seconds) | 0.0075 | |
| **Object Identification** | | |
| Method | IsodataThreshold | |
| Value | -0.924 | |
| **Object Selection Parameter** | **Min** | **Max** |
| ObjectAreaCh1 | 250 | 5136 |
| ObjectShapeP2ACh1 | 1.115 | 6.877 |
| ObjectShapeLWRCh1 | 1 | 10 |
| ObjectAvgIntenCh1 | 10 | 1612 |
| ObjectVarIntenCh1 | 0 | 32767 |
| ObjectTotalIntenCh1 | 0 | 10000000000 |
| **Display Options** | | |
| Composite Color (Hex) | #0000FF | |
| SelectedObject | #0000FF | |
| RejectedObject | #FF7F00 | |
| MaskCh2 | #FF0000 | |

| Channel 2: p53 | | |
|---|---|---|
| Dye | XF93 - TRITC | |
| Apply Illumination Correction | False | |
| Apply Background Correction | True | |
| Gain | 100 | |
| Z Offset | 0.00 | |
| Step Size | 0.00 | |
| Number of Steps | 0 | |
| Projection Method | | |
| Projection Direction | None | |
| Detection Mode | Widefield | |
| Grid Type | | |
| Pin Hole Size | | |
| Intensity Percent | 100 | |
| **Exposure Parameters** | | |
| Method | Fixed | |
| Exposure Time (seconds) | 0.05 | |
| **Object Identification** | | |
| Method | None | |
| Value | 0 | |
| **Object Selection Parameter** | **Min** | **Max** |
| AvgIntenCh2 | 0 | 1000 |
| TotalIntenCh2 | 0 | 10000000000 |
| **Display Options** | | |
| Composite Color (Hex) | #FF0000 | |
| SelectedObject | #0000FF | |
| RejectedObject | #FF7F00 | |
| MaskCh2 | #FF0000 | |

| Assay | |
|---|---|
| Assay Algorithm | TargetActivation.V4 |
| Assay Version | 6.0 (Locally Installed Version: 6.0.0.4004) |
| Focus Channel | 1 |
| #Channels | 2 |
| **Assay Parameters** | |
| PixelSize | 0.645 |
| Type_1_EventDefinition | 0 |
| Type_2_EventDefinition | 0 |
| Type_3_EventDefinition | 0 |
| AvgIntenCh2LevelHigh | 32767 |
| AvgIntenCh2LevelHigh_CC | 2 |
| AvgIntenCh2LevelLow | 34.095 |
| AvgIntenCh2LevelLow_CC | 10 |
| MinRefAvgObjectCountPerField | 2 |
| ObjectAreaCh1LevelHigh | 59.535 |
| ObjectAreaCh1LevelHigh_CC | 2 |
| ObjectAreaCh1LevelLow | 27.783 |
| ObjectAreaCh1LevelLow_CC | 10 |
| ObjectAvgIntenCh1LevelHigh | 630 |
| ObjectAvgIntenCh1LevelHigh_CC | 2 |
| ObjectAvgIntenCh1LevelLow | 305 |
| ObjectAvgIntenCh1LevelLow_CC | 10 |
| ObjectShapeLWRCh1LevelHigh | 1.67 |
| ObjectShapeLWRCh1LevelHigh_CC | 2 |
| ObjectShapeLWRCh1LevelLow | 1.19 |
| ObjectShapeLWRCh1LevelLow_CC | 10 |
| ObjectShapeP2ACh1LevelHigh | 1.26 |
| ObjectShapeP2ACh1LevelHigh_CC | 2 |
| ObjectShapeP2ACh1LevelLow | 1.1 |
| ObjectShapeP2ACh1LevelLow_CC | 10 |
| ObjectTotalIntenCh1LevelHigh | 29767.5 |
| ObjectTotalIntenCh1LevelHigh_CC | 2 |
| ObjectTotalIntenCh1LevelLow | 10835.37 |
| ObjectTotalIntenCh1LevelLow_CC | 10 |
| ObjectVarIntenCh1LevelHigh | 330 |
| ObjectVarIntenCh1LevelHigh_CC | 2 |
| ObjectVarIntenCh1LevelLow | 55 |
| ObjectVarIntenCh1LevelLow_CC | 10 |
| TotalIntenCh2LevelHigh | 1000000000000 |
| TotalIntenCh2LevelHigh_CC | 2 |
| TotalIntenCh2LevelLow | 3311.385 |
| TotalIntenCh2LevelLow_CC | 10 |
| UseMicrometers | 0 |
| VarIntenCh2LevelHigh | 60 |
| VarIntenCh2LevelHigh_CC | 2 |
| VarIntenCh2LevelLow | 0 |
| VarIntenCh2LevelLow_CC | 10 |
| BackgroundCorrectionCh1 | 33 |
| BackgroundCorrectionCh2 | 50 |
| MaskModifierCh2 | 0 |
| NucCleanupCh1 | 1 |
| ObjectSegmentationCh1 | 10 |
| ObjectSmoothFactorCh1 | 1 |
| ObjectTypeCh1 | 0 |
| RejectBorderObjectsCh1 | 1 |
| UseReferenceWells | 0 |

**Well Feature Extents**

| Feature Name | Lower Extent | Upper Extent |
|---|---|---|
| ValidObjectCount* | 0 | 100 |
| SelectedObjectCount | 20 | 750 |
| %SelectedObjects | 0 | 100 |
| ValidFieldCount | 0 | 111 |
| SelectedObjectCountPerValidField | 0 | 89.9708044982699 |
| EventType1ObjectCount | 0 | 100 |
| %EventType1Objects | 5 | 15 |
| EventType2ObjectCount | 0 | 100 |
| %EventType2Objects | 0 | 100 |
| EventType3ObjectCount | 0 | 100 |
| %EventType3Objects | 0 | 100 |
| MEAN_ObjectAreaCh1 | 0 | 10000000 |
| SD_ObjectAreaCh1 | 0 | 1048576 |
| SE_ObjectAreaCh1 | 0 | 100 |
| CV_ObjectAreaCh1 | 0 | 100 |
| %RESPONDER_ObjectAreaCh1 | 0 | 100 |
| MEAN_ObjectShapeP2ACh1 | 0 | 100 |
| SD_ObjectShapeP2ACh1 | 0 | 100 |
| SE_ObjectShapeP2ACh1 | 0 | 100 |
| CV_ObjectShapeP2ACh1 | 0 | 100 |
| %RESPONDER_ObjectShapeP2ACh1 | 0 | 100 |
| MEAN_ObjectShapeLWRCh1 | 0 | 100 |
| SD_ObjectShapeLWRCh1 | 0 | 100 |
| SE_ObjectShapeLWRCh1 | 0 | 100 |
| CV_ObjectShapeLWRCh1 | 0 | 100 |
| %RESPONDER_ObjectShapeLWRCh1 | 0 | 100 |
| MEAN_ObjectTotalIntenCh1 | 0 | 10000000000 |
| SD_ObjectTotalIntenCh1 | 0 | 10000000000 |
| SE_ObjectTotalIntenCh1 | 0 | 100 |
| CV_ObjectTotalIntenCh1 | 0 | 100 |
| %RESPONDER_ObjectTotalIntenCh1 | 0 | 100 |
| MEAN_ObjectAvgIntenCh1 | 0 | 4095 |
| SD_ObjectAvgIntenCh1 | 0 | 4095 |
| SE_ObjectAvgIntenCh1 | 0 | 100 |
| CV_ObjectAvgIntenCh1 | 0 | 100 |
| %RESPONDER_ObjectAvgIntenCh1 | 0 | 100 |
| MEAN_ObjectVarIntenCh1 | 0 | 10000000 |
| SD_ObjectVarIntenCh1 | 0 | 10000000 |
| SE_ObjectVarIntenCh1 | 0 | 100 |
| CV_ObjectVarIntenCh1 | 0 | 100 |
| %RESPONDER_ObjectVarIntenCh1 | 0 | 100 |
| MEAN_TotalIntenCh2 | 0 | 10000000000 |
| SD_TotalIntenCh2 | 0 | 10000000000 |
| SE_TotalIntenCh2 | 0 | 100 |
| CV_TotalIntenCh2 | 0 | 100 |
| %RESPONDER_TotalIntenCh2 | 0 | 100 |
| MEAN_AvgIntenCh2 | 0 | 4095 |
| SD_AvgIntenCh2 | 0 | 4095 |
| SE_AvgIntenCh2 | 0 | 100 |
| CV_AvgIntenCh2 | 0 | 100 |
| %RESPONDER_AvgIntenCh2 | 0 | 100 |
| MEAN_VarIntenCh2 | 0 | 10000000 |
| SD_VarIntenCh2 | 0 | 10000000 |
| SE_VarIntenCh2 | 0 | 100 |
| CV_VarIntenCh2 | 0 | 100 |
| %RESPONDER_VarIntenCh2 | 0 | 100 |

* Indicates the default well feature for the Assay Protocol
** Indicates feature extents are dependent upon system reference well settings

| Selected Well Features to Store |
|---|
| Status |
| TargetActivationV4Well:MEAN_AvgIntenCh2 |
| TargetActivationV4Well:MEAN_ObjectAreaCh1 |
| TargetActivationV4Well:MEAN_ObjectAvgIntenCh1 |
| TargetActivationV4Well:MEAN_ObjectShapeLWRCh1 |
| TargetActivationV4Well:MEAN_ObjectShapeP2ACh1 |
| TargetActivationV4Well:MEAN_ObjectTotalIntenCh1 |
| TargetActivationV4Well:MEAN_TotalIntenCh2 |
| TargetActivationV4Well:SelectedObjectCount |
| TargetActivationV4Well:SelectedObjectCountPerValidField |
| TargetActivationV4Well:ValidFieldCount |
| TargetActivationV4Well:ValidObjectCount |

| Selected Cell Features to Store |
|---|
| TargetActivationV4Cell:AvgIntenCh2 |
| TargetActivationV4Cell:Cell# |
| TargetActivationV4Cell:Height |
| TargetActivationV4Cell:Left |
| TargetActivationV4Cell:ObjectAreaCh1 |
| TargetActivationV4Cell:ObjectShapeLWRCh1 |
| TargetActivationV4Cell:ObjectShapeP2ACh1 |
| TargetActivationV4Cell:ObjectTotalIntenCh1 |
| TargetActivationV4Cell:Top |
| TargetActivationV4Cell:TotalIntenCh2 |
| TargetActivationV4Cell:Width |

#### Analysis Sequence "siRNA_Screen_TP53_analysis_May2020" (Opera Phenix)

**Microscope Acquisition Settings**
20X Air, Non-confocal, Binning: 2
DAPI: 120ms exposure, 100% power, height 2.0um
A488: 100ms exposure, 100% power, height 2.0um
9 fields per well, no stack

| Step | Input | Method | Output |
|---|---|---|---|
| **Input Image** | **Flatfield Correction :** None Brightfield Correction<br>**Stack Processing :** Individual Planes<br>**Min. Global Binning :** Dynamic | | |
| **Find Nuclei** | **Channel :** DAPI<br>**ROI :** None | **Method :** B<br>Common Threshold : 0.35<br>Area : > 70 µm² Splitting<br>Coefficient : 7 Individual<br>Threshold : 0.15<br>Contrast : > 0.18 | Output Population : Nuclei |
| **Select Population** | **Population :** Nuclei | **Method :** Common Filters<br>Remove Border Objects<br>Region : Nucleus | Output Population : Nuclei Valid |
| **Select Cell Region** | **Population :** Nuclei Valid | **Method :** Resize Region [%]<br>Region Type : Ring Region<br>Outer Border : -50 %Inner Border : 0 % | Output Region : Perinuclear Ring Region |
| **Calculate Intensity Properties** | **Channel :** Alexa 488<br>**Population :** Nuclei Valid<br>**Region :** Nucleus | **Method :** Standard Mean Contrast | Property Prefix : TP53 Mean Intensity Nucleus Alexa 488 |

| Calculate Intensity Properties (2) | Input | Method | Output |
| --- | --- | --- | --- |
| | **Channel :** Alexa 488<br>**Population :** Nuclei Valid<br>**Region :** Perinuclear Ring Region | **Method :** Standard Mean | Property Prefix : TP53 Mean Intensity Perinuclear Ring Region Alexa 488 |

| Calculate Properties | Input | Method | Output |
| --- | --- | --- | --- |
| | **Population :** Nuclei Valid | **Method :** By Formula<br>Formula : A/B Variable A : TP53 Mean Intensity Nucleus Alexa 488 Mean Variable B : TP53 Mean Intensity Perinuclear Ring Region Alexa 488 Mean | Output Property : TP53 Mean intensity Nu/cyto |

| Define Results | Results |
| --- | --- |
| | **Method :** List of Outputs<br>**Population : Nuclei Valid**<br>Number of Objects<br>TP53 Mean Intensity Nucleus Alexa 488 Mean : Mean<br>TP53 Mean Intensity Nucleus Alexa 488 Contrast : Mean<br>TP53 Mean Intensity Perinuclear Ring Region Alexa 488 Mean : Mean TP53 Mean intensity Nu/cyto : Mean<br><br>**Object Results**<br>Population : Nuclei : None<br>Population : Nuclei Valid : None |

Acapella version: 5.0.1.124082. Timestamp: 2020-05-06 16:11:24 +1000.
