## Supplementary Information (including Methods and Supplementary Figures) for "Nuclear stabilisation of p53 requires a functional nucleolar surveillance pathway"

### Supplementary Materials and Methods

**Cell Lines and Cell Culture:** A549 (human lung adenocarcinoma, CCL-185) and HEK-293 (CRL-1573) cell lines were obtained from the ATCC. The Flp-In™ T-Rex™ U2OS cells[1] were a generous gift from Prof. Laurence Pelletier (Lunenfeld-Tanenbaum Research Institute, Toronto, Canada). Unless otherwise stated, cell culture reagents were purchased from Gibco (ThermoFisher Scientific). All cell lines were cultured at 37°C with 5% $CO_2$ in complete growth media. A549 cells were cultured in Dulbecco's Modified Eagle Medium: Nutrient Mixture F-12 (DMEM:F12) with HEPES, supplemented with 10% foetal bovine serum (JRH Biosciences #12003C or Sigma-Aldrich #F9423) and 2 mM GlutaMAX™ or 2mM L-Glutamine. HEK-293 cells were cultured in DMEM supplemented with 10% FBS (Sigma-Aldrich #F9423) and 2mM L-glutamine. The Flp-In™ T-Rex™ U2OS cells were cultured in DMEM supplemented with 10% FBS and 1 μg/mL Blasticidin (Melford). For expression of MDM2 protein, the cDNA for the human full-length MDM2 sequence was cloned into the pcDNA5 vector (ThermoFisher Scientific) to enable expression of the proteins with an N-terminal 2xFLAG-PreScission protease site-His6 (FLAG) tag. The plasmid was then transfected into the cells for stable integration into the host genome, and selected using Hygromycin B (Formedium) according to the manufacturers' instructions.

**Generation of Mouse Embryonic Fibroblasts (MEFs):** All animal experiments were performed with approval from the Australian National University Animal Experimentation Ethics Committee (protocol numbers A2015/58, A2018/56). To generate MEFs, C57BL/6 Mdm2$^{C305F/C305F}$ mice (a gift from Prof Yanping Zhang, University of North Carolina, Chapel Hill, USA) or their wild-type C57BL/6 littermates were plug mated. Mouse embryos were isolated at E13.5 and prepared as similarly described[2], digesting individual embryos in 1 mL of 0.25% trypsin-EDTA (Gibco) containing 100U DNaseI (Sigma-Aldrich #D5025) at 37°C at 10% $CO_2$ for 15 minutes. To stop the digestion, 1 mL of DMEM (Gibco) supplemented with 10% FBS (Sigma-Aldrich #F9423), 1x Antibiotic-Antimycotic, 2 mM GlutaMAX™ and 1x MEM non-essential amino acids (NEAA) was added, and the cells were centrifuged at 300 x g for

5 minutes to pellet the cells. Cell pellets were then resuspended and cultured in fresh DMEM containing 10% FBS, 1x Antibiotic-Antimycotic, 2 mM GlutaMAX$^{TM}$ and 1x MEM NEAA at 37$^{o}$C.

**Antibodies:** The α-p53 (human, DO-1, sc-126) antibody was obtained from Santa Cruz Biotechnology. The α-p21 (12D1, #2947S), α-p53 (mouse, 1C12, #2524S) and α-Myc-tag (9B11, #2276) were purchased from Cell Signalling Technologies. The α-RPL5 (ab137617), α-RPL11 (ab79352), α-RPS18 (ab91293), α-RPL21 (ab215724), α-RPS19 (ab57643), α-RPL28 (ab193164) and α-RPL22 (ab77720) antibodies were procured from Abcam. The α-BrDU (B44, #347580) was obtained from BD Biosciences. The α-Actin antibody (C4, #8691001) was acquired from MP Biomedicals, while the α-FLAG (F-3165) antibody was purchased from Merck Millipore. The goat α-mouse (#1706516) and α-rabbit (1706515) HRP-conjugated IgG (H+L) antibodies were obtained from BioRad. Alexa-Fluor 594 donkey α-mouse IgG (H+L, A21203) and Alexa-Fluor 488 goat α-mouse IgG (H+L, A11001) were purchased from ThermoFisher Scientific. Fluorescein isothiocyanate (FITC) conjugated sheep α-mouse IgG (Cappel, ICN Biomedical #55516) and horse anti-mouse IgG (FI-2000, Vector Labs) were also purchased.

**Chemicals/Buffers/General Reagents:** Actinomycin D (A9415), α-Amanitin (A2263), Bovine Serum Albumin (BSA), 5-bromo-2′-deoxyuridine (BrdU, B5002), Cycloheximide, DAPI (4',6-diamidino-2-phenylindole), Diethyl pyrocarbonate (DEPC), Dimethyl Sulfoxide (DMSO), Doxorubicin (D1515), 5-Fluorouracil (F6627), Leptomycin B (L2913), MG132 (C2211), Propidium iodide (PI), Sucrose, Tergitol/IGEPAL (NP-40), Triton™ X-100, Tween®20 were purchased from Sigma-Aldrich. Camptothecin (S1288) and Etoposide (S1225) were purchased from Selleck Chemicals and CX-5461 (SYN-3031) was obtained from SYNkinase. Dulbecco’s phosphate-buffered saline (dPBS, pH 7.4, Gibco) and ultrapure water (Invitrogen) were procured from ThermoFisher Scientific. 5X siRNA buffer (B-002000-UB-100, diluted to 1X with ultrapure water), DharmaFECT1 transfection reagent (T-2001) and assay-specific siRNAs (further details provided below) were purchased from Horizon Discovery. Polyethylenimine (PEI, #23966) was purchased from Polysciences. Paraformaldehyde (PFA, diluted in dPBS prior to use) was obtained from Electron Microscopy Sciences (Fisher Scientific). Complete protease inhibitor cocktail and PhoSTOP phosphatase inhibitor were purchased from Roche. All compounds used for treatment of cells, apart from α-Amanitin ($H_2O$), CX-5461 (50 mM $NaH_2PO_4$, pH 4.5) and Leptomycin B (70% methanol) were solubilised in DMSO and stored at -20$^{o}$C in aliquots prior to use.

**siRNA transfection:** For candidate-based studies, A549 cells were reverse transfected with up to 50 nM (final concentration) of candidate-specific siRNA (Horizon Discovery, please see **Supplementary Files 1** and **6** for catalogue numbers for each specific candidate) or non-targeting siRNA (ONTARGETplus (OTP) non-targeting control siRNA, Horizon Discovery, D-001810-10-50, prepared as per manufacturers' instructions). Briefly, transfection mixes containing DMEM:F12 basal media with DharmaFECT1 (T-2001, final concentration of 1µL/mL) were prepared with siRNA and allowed to complex for 20 minutes. A549 cells (prepared in suspension in DMEM:F12 growth media) were then added on top and the cells incubated at 37$^{o}$C in 5% $CO_2$. At 24 hours' post transfection, media was removed and replaced with fresh growth media. For the transfection of U2OS Flp-In cells, siRNAs (final concentration of 50 nM) were prepared by complexing with Lipofectamine$^{TM}$ RNAiMAX transfection reagent (in Opti-MEM media, ThermoFisher Scientific) prior to reverse transfection of cells (as similarly described above).

**siRNA screening:** The primary genome-wide siRNA screening was performed using the Dharmacon Human siGENOME SMARTpool siRNA library (Horizon Discovery, G-005005-E2); the secondary screen using individual duplex siRNA was performed using the Human siGENOME siRNA library (set of 4 duplexes, Horizon Discovery, GU-005005-E2). For the custom libraries, human siGENOME SMARTpool siRNAs were cherrypicked for individual genes (catalogue details available in **Supplementary Files 1 and 6**). Transfection mixes containing DMEM:F12 basal media, DharmaFECT1 (T-2001, final concentration of 1µL/mL; i.e. 0.038 µL in 37.5µL final volume) and siRNA were complexed for 20 minutes in Corning Costar 384 well optical microplates (#3712). We used 40 nM final concentration for the primary screen, 25 nM final concentration for the secondary screen and either 10 nM of non-targeting siRNA (for the "p53 stabilisation" and follow-up co-depletion screens), 10 nM siRPS19 siRNA (for the "modifiers of ribosomal stress" and follow-up co-depletion screens) or 10 nM siHEATR3, siRPL5 or siRPL11 siRNAs (for the follow-up co-depletion screens; details for siRNAs located in **Supplementary Files 1 and 6**). Using a reverse transfection strategy, A549 cells (750 cells/well prepared in DMEM:F12 growth media) were then delivered on top of the transfection mixtures; plates were briefly centrifuged at 500 x g for 1 minute, and then incubated in a LiCONiC microplate incubator at 37$^{o}$C with 5% $CO_2$ for 24 hours. At 24 hours post-transfection, media was removed and replaced with fresh DMEM:F12 growth media and returned to the incubator. At 72 hours post transfection, media was removed and cells were fixed with 4% PFA (25 µl/well) for 10 minutes, washed with dPBS (50 µl/well) and permeabilised with of 0.5% TTX-100 in dPBS (25 µl/well) containing 0.5 µg/mL DAPI for a further 10 minutes. Plates were then washed with dPBS twice (50 µl/well). Cells were immunostained with anti-p53 (DO-1) antibody in a 1% BSA in dPBS solution, washed twice with dPBS, followed by the addition of the Alexa-Fluor secondary antibody for signal detection. After the final dPBS wash, stained cells were left in dPBS (50 µl/well), the plates heat-sealed with foil seals (Agilent PlateLoc) and imaged on either a Cellomics ArrayScan VTi (ThermoFisher Scientific) or a Perkin Elmer Opera Phenix high content imaging microscope. Details for image acquisition and quantitative algorithms can be found in **Supplementary File 8**.

**siRNA screening analysis:** As both primary screens were combinatorial screens (i.e. co-transfecting two siRNAs simultaneously), we included two sets of controls within the plate; the 'health' controls (which were transfected with one siRNA only, to ensure that the cell viability response was consistent within each screen run, and that co-transfection of siRNAs did not impact on cell viability), and the 'normalising' controls for each independent screen (two siRNAs co-transfected, which were also used for quality control and normalisation of data). We also included some 'mock' (lipid-only) controls which contained the single siRNA which was being co-transfected with the unknowns on the plates. The approach for analysis of each screen is outlined below. We also utilised a similar approach for data normalisation for all subsequent screening experiments. For the primary screens, within the library (as provided by the manufacturer), there were 55 duplicate wells; we opted to remove any duplicate candidates from the library which were screened more than once, where we removed the duplicate(s) that were the least changed when compared to the respective normalisation strategy for the screen (therefore leaving 18,120 unique candidates screened). Using RNAseq data generated from transfection of A549 cells with either siOTP-NT ("p53 stabilisation" screen) or siRPS19 siRNA ("modifiers of ribosomal stress" screen) for 72 hours, we removed 'non-expressed' candidates from the screening datasets (based on an RPKM value cut-off of < 0.05). All subsequent analyses were performed on the 'expressed' candidates (13,855 and 14,577) in the "p53 stabilisation" and "modifiers of ribosomal stress" screens, respectively.

*"p53 stabilisation screen" analysis:* Each plate contained multiple wells of three different negative control conditions (siOTP-NT, siTP53 and mock transfected) and one positive control (siRPS19). For the normalisation controls, to balance the absolute amount of siRNA and to replicate precise screening conditions, each control was co-transfected with siOTP-NT (i.e. siOTP-NT+siOTP-NT, siTP53+siOTP-NT, siRPS19+siOTP-NT etc). For the health controls, we normalised to the average siOTP-NT value (presented as a fold-change, FC); for the normalisation controls and library candidates, we normalised to the average of the siOTP-NT+siOTP-NT wells (data presented as FC) – this approach was used for both p53 intensity and cell number outputs and used for evaluating quality control metrics. Our QC metrics also focused on specific control combinations – siOTP-NT (negative control which had some baseline p53 expression) and siRPS19 (which enhanced nuclear p53 stabilisation). We calculated the average health (siOTP-NT only) and normalisation (siOTP-NT+siOTP-NT) controls to be 1.02 ± 0.19 (mean % co-efficient of variation for each library plate screened; %CV = 18.49) and 1.02 ± 0.19 (mean %CV = 18.49), respectively. The positive controls (siRPS19 and siRPS19+siOTP-NT) scored 5.86 ± 0.86 (mean %CV = 14.61) and 6.17 ± 0.85 (mean %CV= 13.68), respectively. The Z'-factor calculated for siOTP-

NT+siOTP-NT vs siRPS19+siOTP-NT for each plate (approach as similarly described[3]) demonstrated a very robust result of 0.4 ± 0.16. A representative scatterplot of normalised p53 intensity values for the "p53 stabilisation" primary screen is presented in **Supplementary Fig. 7a**. All data was normalised to the average siOTP-NT+siOTP-NT value for each plate, ensuring we could directly use our experimentally-determined 2-fold increase in p53 as the cut-off (**Supplementary Fig. 1a-d**) for further analysis. For graphical presentation of the data, FC data was converted to $\log_2$ (p53 FC value).

*"Modifiers of ribosomal stress" screen:* We used a similar approach described above for the health (single-siRNA) controls, and obtained p53 intensity FC values (normalised to siOTP-NT) of 1.02 ± 0.20 (mean %CV across entire screen = 19.39%) for siOTP-NT and 6.14 ± 1.01 (mean %CV = 16.43%) for siRPS19 overall. For this screen, the control combinations for QC were different, because depletion of RPS19 leads to an extremely robust and strong increase in p53 stabilisation. Therefore, the 'normalisation' controls for this screen were the siOTP-NT+siRPS19 (negative control) and the siTP53+siRPS19 (positive control). The data was normalised to the average negative control value (siOTP-NT+siRPS19; 1.01 ± 0.17, mean %CV = 16.63%). The positive control (siTP53+ siRPS19) averaged 0.30 ± 0.06 (mean %CV = 19.47%) across the entire screen. The overall Z' factor was weaker than the 'p53 stabilisation screen' but still significant on average (0.04 ± 0.19) (**Supplementary Fig. 7b**). To identify candidates from this screen, we applied a cut-off of 3SD above the average positive control FC readout (siTP53+siRPS19) (calculated to be 0.48; relaxed to 0.5). Based on these criteria, we analysed 64 candidates using the Dharmacon siGENOME SMARTpool individual duplexes (25 nM/duplex). We observed in some cases that the duplexes demonstrated a weaker suppression of p53 induced by RPS19 depletion compared to the SMARTpool screen (likely because multiple siRNAs targeting the same mRNA transcript more efficiently deplete the target), therefore, we relaxed the hit cut-off criteria to a p53 FC value of ≤ 0.65. We proceeded to further validate candidates that had 2 or more duplexes fit these criteria.

**Alphascreen p53 analysis:** The levels of p53 protein were quantitatively analysed using the Alphascreen SureFire Total p53 Assay Kit (TGR Biosciences/Perkin Elmer #TGRT53). Briefly, A549 cells (40,000 cells/well) were reverse transfected with 40 nM siRNA (final reaction volume 0.5 mL) in Nunc 24-well cell culture plates for 72 hours, refreshing media at 24 hours' post transfection. Just prior to harvest, in order to normalise p53 signal, plates were imaged on the IncuCyte S3 Live-Cell Analysis Imaging System (Essen Biosciences) using the scan-on-demand feature (2015A software release), calculating the Phase Object Confluence (Percent) for each well. Media was then carefully removed and the cell monolayers lysed with 30 µL 1x Alphascreen Lysis Buffer for 2-3 minutes on an orbital rocker, prior to transfer of lysates to -20$^{o}$C. Lysates (4 µL/well in Perkin Elmer 384 well proxiplates, #6008280) were then assayed in as per the manufacturers' instructions. Plates were then read using the Perkin Elmer EnSpire Multimode Plate Reader instrument using the instrument pre-defined Alphascreen settings and the data normalised to the cell confluence readings, then normalised to the non-targeting siRNA treated wells.

**Cell Treatments:** Either A549 cells (0.9 x $10^5$/well, transfected as described above) or MEFs (~3 x $10^5$/well) were plated in Nunc 6-well tissue culture dishes and incubated for 24 hours. For determination of candidate-specific knockdown (where appropriate), unless otherwise described, cells were harvested at 24 hours' post transfection/seeding; otherwise, the transfection/culture media was removed and replenished with either fresh media, or pharmacological inhibitors at the required dose (prepared in growth media), and incubated for the desired period of time (up to 72 hours). For cells which were UV treated, subjected to heat-shock (45$^{o}$C) or irradiated (IR) with gamma irradiation, cells were plated down for 69 hours, treated with 5 mJ/cm$^2$ using a UV Crosslinker (UV Stratalinker 1800), heated at 45$^{o}$C for 30 minutes, or exposed to 10 Gy (Rad Source RS 2000), returned immediately to a 37$^{o}$C incubator with 5% $CO_2$ and incubated for 3 hours prior to harvested for immunoblotting.

**Cell Cycle Analysis:** Cells were pulse labelled with 10 µM BrdU and incubated for 30 minutes at 37$^{o}$C 5% $CO_2$. Media was collected, the cell monolayer washed with dPBS then the adherent cells trypsinised for 5 minutes at room temperature with 0.25% Trypsin-EDTA. Trypsinised cells were then neutralised

(1:1) with growth media. The media, washes and trypsinised cells were pooled, and the cells centrifuged at 1200 rpm for 5 minutes to pellet the cells. Cells were then fixed with ice-cold 80% ethanol. Prior to FACS analysis, cells were permeabilised (1M HCl, 0.5% TTX-100) then incubated with α-BrDU antibody, followed by FITC/Alexa-Fluor-488 conjugated α-mouse IgG secondary antibody. Cells were then stained with PI (0.01 mg/mL) and analysed on either a BD FACS CantoII or LSR instrument. All FACS data was analysed using FlowLogic Software (Inivai Technologies).

**Plasmids and cloning:** FLAG-tagged human RPL5 and RPL11 (FLAG-RPL5, FLAG-RPL11) constructs have been previously described[4]. To generate the human myc-tagged (MT) human HEATR3 construct, the MGC clone containing the full-length human HEATR3 cDNA (MHS6278-202832721) was purchased from GE Life Sciences as a template for PCR cloning. The pcDNA3.1 vector (containing a puromycin selection cassette) was a kind gift from A/Prof Armadeep Dhillon (LaTrobe University, Bundoora, Australia). Primers for cloning were purchased from Sigma-Aldrich. The 5' (forward) primer (5'-GAATTCGGATCCGCCACCATGGGGGAACAAAAACTCATCTCAGAAGAGGATCTGGGCAAGAGCCGGACGAAG-3') was engineered to include a 5' *BamHI* restriction site, immediately proceeded by a nucleotide sequence that, when translated in-frame, generates a methionine residue and the myc-tag (M-EQKLISEEDL), followed by the endogenous human HEATR3 sequence (residues 2-680). The reverse primer (5'-GAATTCCTCGAGTTAAGAAGTCAGTCTTTTCTCAACAGTTTC-3') contained a *XhoI* restriction site immediately after the stop codon. The cDNA template was incubated with the KOD Hot Start Mastermix (Novagen) and 10 µM of forward and reverse primers, and amplified in a BioRad Thermal Cycler using the following conditions (95$^{o}$C for 2 minutes, three cycles at 95$^{o}$C for 20 seconds, annealing at 62$^{o}$C for 10 seconds and extension at 70$^{o}$C for 45 seconds, followed by 33 cycles of denaturation at 95$^{o}$C for 20 seconds, annealing at 67$^{o}$C for 10 seconds and extension at 70$^{o}$C for 45 seconds). The PCR product (2106 bp) was purified using the Nucleospin Gel and PCR Clean-up kit (Macherey Nagel) and along with the pcDNA 3.1 vector, digested at 37$^{o}$C with *BamHI* and *XhoI* restriction enzymes as per the manufacturers' instructions (Promega) for 1.5 hours. Digests were then incubated at 65$^{o}$C for 10 minutes to denature the restriction enzymes, before electrophoresing on 1.2%

Tris-Acetate-EDTA (TAE) agarose gels containing Sybr-Safe (ThermoFisher Scientific) at 100V for ~1.5 hours. The digested insert and vector were then extracted from the gel and cleaned up using the Macherey-Nagel Gel and PCR Clean-Up Kit (as per manufacturers' instructions). The concentration of the DNA was determined using a Nanodrop 2000 instrument (ThermoFisher Scientific), and a ligation reaction prepared as per manufacturers' instructions using the Promega T4 DNA Ligase Kit at a 1:3 (vector: insert) ratio. The reaction was then incubated at $65^{o}$C for 10 minutes to denature the enzyme, and cooled on ice for 2 minutes. The ligation reaction was added to Bioline chemically competent bacteria (BIO-85027), incubated on ice for 30 minutes, heated at $42^{o}$C for 90 seconds and then placed on ice for 2 minutes. Cells were recovered in 1 mL of Super Optimal broth with Catabolite repression (SOC) media (Bioline) for 1 hour at $37^{o}$C shaking at 200 rpm prior to plating (under aseptic conditions) onto L-Broth (LB) Agar plates containing 100 µg/mL ampicillin and incubated for 16 hours at $37^{o}$C. Individual colonies were picked and inoculated into 5 mL LB containing 100 µg/mL ampicillin and incubated, shaking at 200 rpm, for 16 hours at $37^{o}$C. From 2 mL of each culture, plasmid DNA was extracted using the Nucleospin Plasmid Kit (Macherey-Nagel) as per manufacturers' instructions. The MT-HEATR3 constructs were sequenced verified by Sanger sequencing (Micromon Genomics, Monash University, Australia) prior to use.

**RNA isolation and cDNA synthesis:** Unless otherwise described, total RNA was harvested and isolated using the NucleoSpin RNA extraction kit (Macherey-Nagel) according to the manufacturer's instructions. Total RNA was quantified using the Nanodrop 2000C Spectrophotometer (Thermo Scientific) prior to storage at $-80^{o}$C. cDNA was prepared using the Invitrogen Superscript III Kit (ThermoFisher Scientific) with random primers (Promega) as previously described[5].

**Primers and real-time quantitative PCR:** Primer design and quantitative real-time PCR (qPCR) analysis was carried out as similarly described[5]. Where appropriate, gene expression was normalised to housekeeping gene expression (*GAPD*) and fold change determined using the relative quantitation ($2^{-(\Delta\Delta Ct)}$) method[6]. A list of primer sequences used in this publication is located in **Supplementary Table 1**.

**RNAseq analysis:** Total RNA for sequencing was isolated from cells using the RNeasy Mini Kit (Qiagen) according to the manufacturers' instructions, and concentration and quality determined using the Agilent 2100 Bioanalyser. Sequencing libraries were prepared using the TruSeq RNA library preparation kit (Illumina) and sequenced on an Illumina HiSeq2500 (6 samples/lane). The generated 50bp paired-end reads were aligned to the genome using TopHat[7] v2.0.8b with default parameters and the reads counted using HTSeq[8]. The differential expression was calculated utilising the DESeq package[9] in R (version 3.0.2). Absolute gene expression was defined by determining reads per kilobase per million (RPKM) as previously described[10].

**Immunoblotting:** Unless otherwise described, cell monolayers were washed with dPBS then lysed in Western Solubilisation Buffer (20 mM HEPES, 0.5 mM EDTA, and 2% SDS). Protein concentration was determined using BioRad DC Assay as per the manufacturers' instructions. Proteins were separated by SDS-polyacrylamide gel electrophoresis, transferred to PVDF membrane (Immobilon-P, Merck Millipore), blocked with 5% skim milk (prepared with Tris-buffered saline containing 0.1% Tween-20; TBST) at room temperature, and incubated with primary antibody either at room temperature (1-2 hours) or overnight at 4°C. Membranes were washed with TBST, incubated with secondary antibodies for 1 hour, then the protein bands visualised using Clarity Western ECL (BioRad) or Western Lightning ECL (Perkin Elmer) and images either acquired digitally using the Bio-Rad Gel Doc System (with Image Lab Touch Software) or onto Amersham Hyperfilm (Cytivia/GE Healthcare).

**Immunofluorescence:** Cells were seeded and/or transfected in CC2-coated 4-well chamber slides (Thermo Fisher Scientific). Cells were fixed at the desired timepoint with 4% PFA, rinsed with dPBS and permeabilised with 0.5% TTX-100 prepared in dPBS containing 0.5 μg/mL DAPI. Slides were rinsed twice with dPBS and incubated with the primary antibody diluted in 1% BSA prepared in dPBS for 1 hour at room temperature, followed by washing with dPBS twice then incubating with the appropriate AlexaFluor-conjugated secondary antibody for 1 hour. After rinsing with dPBS twice, the chamber was removed and a coverslip with Vectashield mounting medium (Vector Laboratories) was placed over the cells and sealed with clear nail polish. Images were captured using an Olympus BX-51 (widefield, 40X objective) using SPOT Advanced software (v4.6, Diagnostic Instruments) and analysed/pseudocoloured images using NIH ImageJ (v2.1.4.6)[11,12] and Adobe Photoshop (CS6).

**Ribosome assembly analysis and p53 ribosomal mapping:** Information about where specific ribosomes integrate into the eukaryotic ribosome was obtained from a paper published by de la Cruz and colleagues[13]. We rescreened the majority of RP genes in a boutique siRNA screen (as similarly described above for the primary screen analysis, but encompassed several RPs which were not screened in the primary screen; details in **Supplementary File 5**), and further stratified the times at which the specific RP was integrated into the ribosome into 6 groups: Early Nucleolar and Nucleolar (Early Nucleolar/Nucleolar), Medium Nucleolar, Late Nucleolar (Medium/Late Nucleolar), Early Nucleolar – Late Positioning, Late Nucleolar – Cytoplasmic, Cytoplasmic and Unknown (**Supplementary File 5**). The p53 intensity data for each RP was then binned according to its assembly group. For mapping of the p53 values for RPs onto the ribosome, we imported the existing 80S human ribosome structure published by Khatter and colleagues (PDB reference 4UGO)[14] into the Pymol Molecular Graphics System (Schrödinger, LLC) and utilised Pymol colours in the red and grey spectrums (ranging from the highest p53 value “firebrick” to the lowest value “grey90”) to manually colour each RP (using data in **Supplementary File 5**) within the ribosome structure. Note that RPLP0, RPLP1, RPLP2 and RPLP12 are missing from the structure (and are therefore not included), and RPSA and RACK1 were not rescreened for this analysis.

**Co-immunoprecipitation of 5S rRNA with MDM2:** U2OS cells expressing FLAG-MDM2 were subjected to siRNA-mediated knockdown of HEATR3 for 48 hours, then treated with 1mg/ml tetracycline (Duchefa Biochemie) for 16 hours to induce MDM2 protein expression. Cells were then harvested by sonication in gradient buffer E (20 mM HEPES pH 8.0, 150 mM KCl, 0.5 mM EDTA, 0.1 mM DTT, 5% glycerol) and loaded onto a 10-40% glycerol gradient containing gradient buffer (20 mM

HEPES pH 8.0, 150 mM KCl, 1.5 mM $MgCl_2$, 1 mM DTT, 0.2% Triton X-100). The gradients were centrifuged in a swTi60 rotor (Beckman L7-80) at 4$^o$C at 52,000 rpm for 1.5 hours. The gradient was then manually fractionated into 200 µl fractions. The non-ribosomal (free) fractions at the top of the gradient (fractions 1-6) were then pooled and immunoprecipitated with either control IgG Sepharose beads or anti-FLAG antibody beads (Sigma-Aldrich) using gradient buffer containing 10 % glycerol. Co-purified RNA was extracted using phenol/chloroform/IAA, ethanol precipitated, separated on an 8% acrylamide / 7M urea gel, transferred to a Hybond-N membrane (GE-Healthcare) and then analysed by northern blotting using a radiolabelled probe (5'-CCGAGATCAGACGAGATCGGGCGCGTTCAGGGTGGTATGG-3') hybridizing to the 5S rRNA (as previously described[4]).

**Co-immunoprecipitation (CoIP) of FLAG-tagged RPL5/11 with myc-tagged HEATR3:** HEK293 cells (200,000/well) were seeded into Nunc 6-well tissue culture plates and incubated overnight at 37$^o$C in 5% $CO_2$. The following day, a total of 0.5 µg of plasmid DNA per well was complexed with PEI (1:4.5 ratio of plasmid DNA:PEI) in DMEM basal media for 20 minutes, prior to addition to the cells (added dropwise to the media). Cells were then incubated at 37$^o$C in 5% $CO_2$ for 48 hours. Transfected cells were washed once with dPBS, and then harvested in CoIP buffer (100 mM Tris pH 8.0, 100 mM NaCl, 1% NP40, Complete protease inhibitor cocktail and PhoSTOP), swirling gently on an orbital rocker for 1 hour at 4$^o$C. Cell lysates were then collected and centrifuged at 14,000 rpm for 10 minutes at 4$^o$C; the supernatant was then collected and the concentration determined using the BioRad DC assay as per manufacturers' instruction. Each CoIP sample (300 ng of protein) was precleared using 10 µL of Novex Protein G Dynabeads (Life Technologies, #10004D) and rotated at 4$^o$C for 1 hour. The pre-cleared lysate was then transferred to eppendorf tubes containing 3 µg anti-FLAG antibody diluted in 1x CoIP buffer and incubated overnight, rotating at 4$^o$C. Dynabeads were then washed four times with 1 mL of ice-cold CoIP buffer. The sample was eluted off the beads by boiling at 95$^o$C for 5 minutes in 25 µL of 2x SDS-PAGE loading buffer (130 mM Tris-HCl, pH 6.8, 0.67% (w/v) SDS, 16% (v/v) glycerol, 0.083% (w/v) bromophenol blue and 19.3 mM β-mercaptoethanol), electrophoresed on 4-20% Tris-Glycine gradient gels (Invitrogen) and immunoblotted as described above.

**Ribosomal subunit and polysome fractionation:** A549 cells (transfected with target-specific siRNAs as described above for 72 hours in 10 $cm^2$ dishes, seeded at 3.2 $x10^5$ cells/plate) were treated with 100 µg/mL cycloheximide for 5 minutes then harvested in hypotonic lysis buffer as similarly described[15]. Lysates from equal cell number (8 $x10^6$) were loaded onto high salt (250 mM NaCl, 3.1-30.1%) or low salt (80 mM NaCl, 10-40% (w/v)) sucrose gradients generated using a BioComp Gradient Master 108, and separated by centrifugation (SW41 rotor at 40,000 rpm for 4 hours or 36,000 rpm for 2.15 hours, respectively) using a Beckmann Coulter Optima XE-100 Ultracentrifuge. Samples were fractionated (1mL fractions) using a Teledyne ISCO Foxy R1 instrument. Absorbance at 260nm was determined using a Brandel UA-6 UV/Vis detector.

**HEATR3 modelling:** An initial alignment of human HEATR3 with *Cheatomium thermophilum* Syo1 (ctSyo1) was extracted from an HHPRED search[16,17]. The ctSyo1 crystal structures 4GMO (free) and 5AFF (in complex with RPL5/11) scored E-values <e-75 over the entire protein sequence despite low sequence identity. To refine the assignment, a consensus secondary structure prediction of HEATR3 was created using PSIPRED4[18] and Quick2D[16]. Subsequently, ctSyo1 secondary structure features (helices of over three residues in length) were extracted from the crystal structures and manually aligned with HEATR3 predictions while maintaining primary sequence similarities using the program ALINE[19]. Additionally, high-confidence prediction of disordered regions[20] with a DISOPRED 3 score higher than 0.5 (over a ten residue stretch) are indicated in grey in the figure. **Supplementary Figure 5** displays the resulting alignment with (coloured residue similarity groups in ALINE with 0.75 cut-off). The manual alignment based on secondary structure-predictions was used to generate a 3D model of HEATR3 with the MODELLER 9.22 suite[21] using PDB 4GMO and 5AFF as templates. The obtained models only diverged in the contraction state between N- and C-terminal regions of the chain, as observed between free and RPL5/11 bound ctSyo1. Positions of RPL11 and the conserved RPL5 N-terminus were modelled based on their interaction with ctSyo1 (PDB 5AFF)[22]. Figures were prepared using the Pymol Molecular Graphics System (Version 2.0, Schrödinger, LLC, https://pymol.org/2/).

**STRING Analysis:** Biological networks were identified from the screening dataset using STRING (versions 9-11) software (http://string-db.org)[23,24], a publicly available online database of physical and functional protein interactions, using the 'high confidence interactions' setting and the KEGG network enrichment analysis feature. Given its large size, in order to organise the network relative to pathways and functional annotation, data obtained from STRING from the "p53 stabilisation" primary screen (**Supplementary File 2**) with an RPKM value of ≥ 0.05 and fold change (FC) p53 value of ≥ 2.0 was plotted using Cytoscape (www.cytoscape.org/), an open source platform for complex network analysis[25] after applying a "force directed layout".

**Gene Ontology and Gene Set Enrichment Analysis:** A gene ontology (GO) enrichment analysis was performed for the genes identified in the "p53 stabilisation screen" with an RPKM ≥ 0.05 and FC (p53) ≥ 2.0 (**Supplementary File 1**) as the query genes, versus all genes from the dataset (as the gene universe) using the ClusterProfiler package[26] in R[27]. We considered three main ontologies: "biological process" (BP), "cellular component" (CC) and "molecular function" (MF); the enrichment *p*-values are corrected for multiple hypothesis testing using the Benjamini-Hochberg procedure within the package, and all the GO terms that were further considered were statistically significantly enriched at the $\alpha$=0.01 level. We then further simplified the GO terms per ontology by removing semantic redundancies of terms using the default method (proposed by Wang and colleagues[28]) which uses the topology of the GO graph structure to compute semantic similarity. We then visualised the top 20 GO terms per ontology across the BP, CC and MF groups (BP is demonstrated in **Fig. 1d** of this manuscript). For the Gene Set Enrichment Analysis (GSEA), we weighted genes based on their FC (p53) values, with a weighting exponent of $p$=1; we limited gene sets based on their size (i.e. $10 \leq$ size of $S \leq 500$), and then selected a statistical significance level of FDR $\leq \alpha$=0.25. The FDR cut-off is consistent with values which have been previously described[29]. From there, we then plotted the GSEA normalised enrichment scores (NES) vs. the GO analysis-based $\log_{10}$-transformed odds-ratios (OR) in a scatterplot (**Supplementary Fig. 2b**) to demonstrate the correlation between both approaches. The Pearson's product moment correlation coefficient between the NES and $\log_{10}$(OR) is r= 0.7512.

**Subcellular Localisation (LOCATE) Analysis:** A cell localisation enrichment analysis was performed for the candidates identified in the "p53 stabilisation screen" with an RPKM $\geq 0.05$ and FC (p53) $\geq 2.0$ (**Supplementary File 4**) as the query genes, versus all genes from the dataset (gene universe) using the LOCATE subcellular localisation database[30], which lists the subcellular localisation of 64,637 human proteins (https://web.archive.org/web/20171231015119/http://locate.imb.uq.edu.au/). The full human database was imported into R software, then converted and parsed to store as an R-readable object. From there, we extracted the subcellular localisation information for all query genes and genes from the gene universe, and performed an enrichment analysis of the number of query genes in every subcellular localisation category. The results were then visualised using a dotplot, where the size of the dot indicates the number query genes found in the localisation category. The $\log_{10}$ odds-ratio (OR) reflects the amount of enrichment/under representation, i.e. $\log_{10}$ OR $< 0$ indicates under-representation, and $\log_{10}$ OR $> 0$ indicates over-representation of query genes in the corresponding category. The coloured bars represent the 95% confidence intervals.

**Statistical Analysis:** Unless otherwise described, statistical tests were performed using GraphPad Prism Software (https://www.graphpad.com) with the relevant test described for each analysis at the appropriate location in the text.

**Supplementary Table 1: Primer Sequences**

| Gene Symbol | Entrez Gene ID | Forward (5'-3') | Reverse (5'-3') | RefSeq |
|---|---|---|---|---|
| FAU | 2197 | CCGTTCAGTCGCCAATATGC | GACTTGATCTTCCGGGGCAA | NM_001997.4 |
| GAPD | 2597 | GGACTCATGACCACAGTCCATGCC | ATGACCTTGCCCACAGCCTTGG | NM_002046.3 |
| RPL5 | 6125 | CAGCGTATGCACACGAACTG | ACCTATTGAGAAGCCTGCGG | NM_000969.3 |
| RPL6 | 6128 | GCACGTGAGAAAACTGCGAG | TTGAGGACCAGAGGTCCAGT | NM_000970.3 |
| RPL10 | 6134 | ACTGAAGATCCTGGTGTCGC | TTCCGCCCCAGGTCAAAAAT | NM_006013.4 |
| RPL11 | 6135 | CTCCATCATGGCGCAGGATCA | GTCTCCACTCTCCCCAACAC | NM_000975.3 |
| RPL12 | 6136 | GGAAGGGCCTGAGGATTACAG | CCCACTGTGTTTAATGTTTTTCTG | NM_000976.3 |
| RPL21 | 6144 | CCATCTTCCAGTAATTCGCCA | CGCATATATGTGGCCAAAGGA | NM_000982.3 |
| RPL22 | 6146 | GCTGCCAATTTTGAGCAGTTT | AGAAAGGCACCTCGGATGTC | NM_000983.3 |
| RPL24 | 6152 | TGTCGCCATGAAGGTCGAG | TCCTCTTGGAAAGGAAAGCCG | NM_000986.3 |
| RPL26 | 6154 | CTCGCGAGATCTTTGGTAAACT | TAAACTTCATTTTGGCCGCTCC | NM_000987.3 |
| RPL28 | 6158 | CGTGGTGGTCATTAAGCGGA | GCCATGCGCAGGTCGG | NM_000991.4 |
| RPL29 | 6159 | TGACCCTATTTCCCGTGCTG | TGCCATTTTCGGGACTGGTT | NM_000992.2 |
| RPL35A | 6165 | CCTCGTCCTTCTCTTACCGC | GAGACCCCGCTTATAGCCAG | NM_000996.2 |
| RPL41 | 6171 | GAAACCTCTGCGCCATGAGA | CTCCACGGTGCAACAAGCTA | NM_021104.1 |
| RPL36A | 6173 | TTGTGCTAAGGCTTGAGTGC | CTGGATCACTTGGCCCTTTCT | NM_021029.5 |
| RPLP1 | 6176 | TCACGGAGGATAAGATCAATGCC | CATTGCAGATGAGGCTCCCA | NM_001003.2 |
| RPLP2 | 6181 | GACGTCATTGCCCAGGGTAT | TCTCATCTTTCTTCTCCTCTGCT | NM_001004.3 |
| RPS4Y1 | 6192 | CAGATTCTCTTCCGTCGCAGA | GTCGATGGACGAGGTGCAAA | NM_001008.3 |
| RPS6 | 6194 | GTGGACGATGAACGCAAACT | TCGGACCACATAACCCTTCCA | NM_001010 |
| RPS7 | 6201 | CCGGATTTTGACGTGCTCTC | ATCTTGGCGCTCGAACTGAA | NM_001011.3 |
| RPS8 | 6202 | CATCTCTCGGGACAACTGGC | CCAATCTTGGTGTTGGCAGC | NM_001012.1 |
| RPS10 | 6204 | CCGCAGAGATGTTGATGCCT | GCACATTCTTGTCTGCCAGC | NM_001014.4 |
| RPS12 | 6206 | AAGCGCCAAGCCCATCTTTG | CAAAGGCCTACCCATTCTCCT | NM_001016.3 |
| RPS14 | 6208 | GAGACGACGTGCAGAAATGG | ATGCAAAGATATGGCAGACACC | NM_005617 |
| RPS17 | 6218 | CCAAAACCGTGAAGAAGGCG | AACCTGCTATCTTGTTGCGG | NM_001021.4 |
| RPS18 | 6222 | GCAGCCATGTCTCTAGTGATCC | GAGCATATCTTCGGCCCACA | NM_022551.2 |
| RPS19 | 6223 | CTCTCGCGAGCTTTCGGAAC | ACAGTAACTCCAGGCATCGTG | NM_001022.3 |
| RPS24 | 6229 | GAGGAAACAAATGGTCATTG | CCAAATACAAAGATGACATCC | NM_001026.4 |
| RPS25 | 6230 | CTCCGAGCTTCGCAATGCC | CCCGAACTTTGCCTTTGGAC | NM_001028.2 |
| RPS26 | 6231 | GCCCGTCTCCTAAGGATTCTC | TTTGTCATCTTGGAGGCACG | NM_001029.3 |
| RPS27 | 6232 | CTACGCACACGAGAACATGC | TCCTGGGCATTTCACATCCAT | NM_001030.4 |
| RXRA | 6256 | TTCCTGCCGCTCGATTTCTC | CAGGGTGCTGATGGGAGAATG | NM_002957 |
| TP53 | 7157 | AAGTCTAGAGCCACCGTCCA | CAGTCTGGCTGCCAATCCA | NM_000546.5 |
| UBA52 | 7311 | AGCTGACGCAAACATGCAGA | CTGCTGGTCAGGTGGGATAC | NM_003333.3 |
| RPL14 | 9045 | TTGGTCGATGGACCTTGCAC | GGACATACTTCTGGTGGGCA | NM_003973.4 |
| HEATR3 | 55027 | TCAAGCGACCTCAGTTCTCC | CTCGGGTGCTGGAGCTTTTC | NM_182922.2 |
| CIRH1A | 84916 | GCTGGAATGGGTGAATTTAAGGT | CATCTGTTCGTGAAACAGCCA | NM_032830.2 |
| RPS4Y2 | 140032 | AAGGAGAGGTTCTGTTCCGT | GATGGACGAGGTGCAAATACA | NM_001039567.2 |

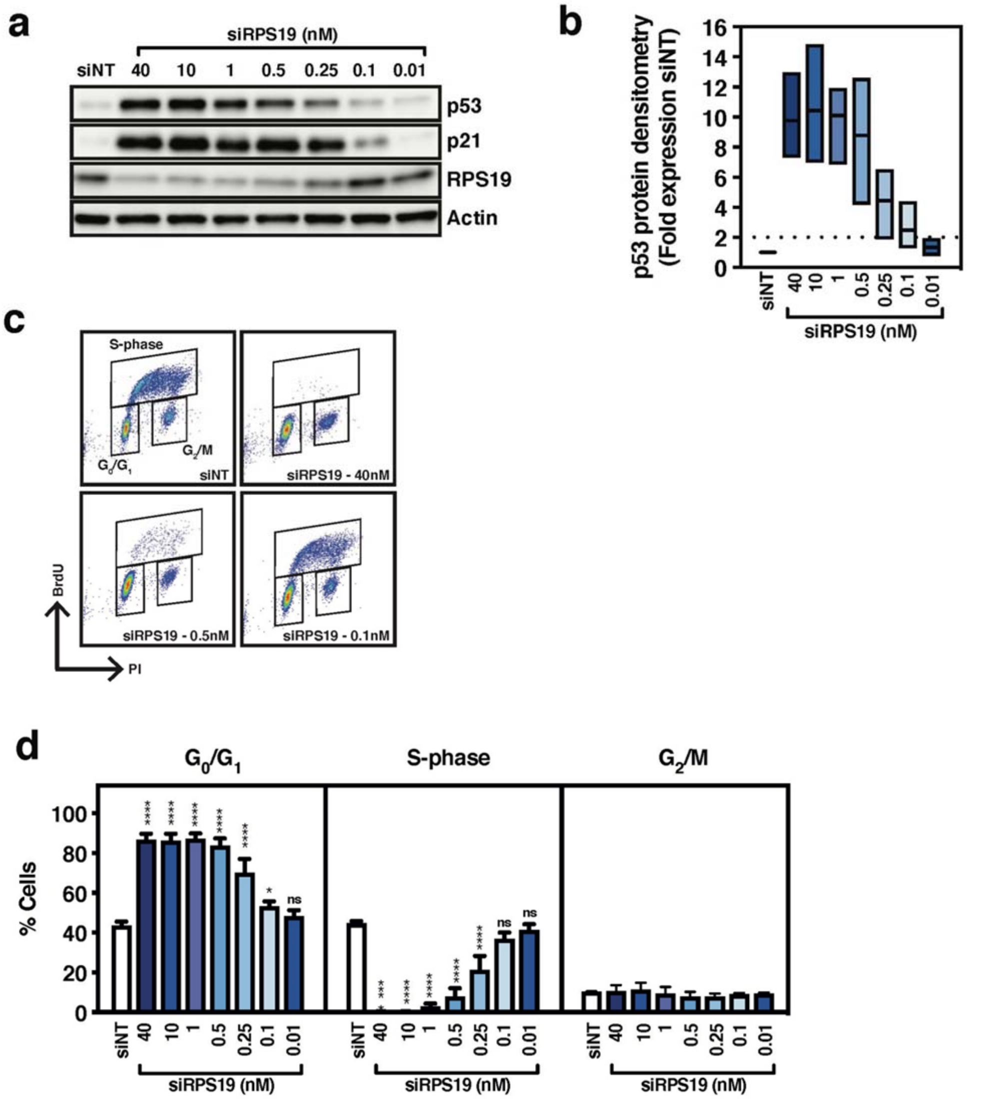


**Supplementary Figure 1: Identifying the minimal level of p53 accumulation sufficient to cause a significant p53-dependent proliferation defect.** To identify the minimum amount of p53 accumulation required to cause a phenotype in cells, we transfected A549 cells for 72 hours with different concentrations of RPS19 siRNA (siRPS19: from 40 – 0.01 nM) and measured p53, p21 and RPS19 protein expression using western blot analysis (**a**), which was subsequently quantified using densitometry analysis (approach outlined in Methods), normalised to non-targeting (siNT) siRNA transfected cells and graphed as mean +/- SD of n=3 (**b**). A549 cells were transfected with the same range of concentrations of siRPS19 (40 – 0.01 nM) for 72 hours and subjected to cell cycle analysis (representative of cell cycle profile is illustrated in **c**, overall analysis graphed as mean +/- SD in **d**). n=3-4 biological experiments, one-way ANOVA with Dunnett's multiple comparison test, ****$p < 0.0001$, *** $p < 0.001$, *$p < 0.05$, ns = not significant.

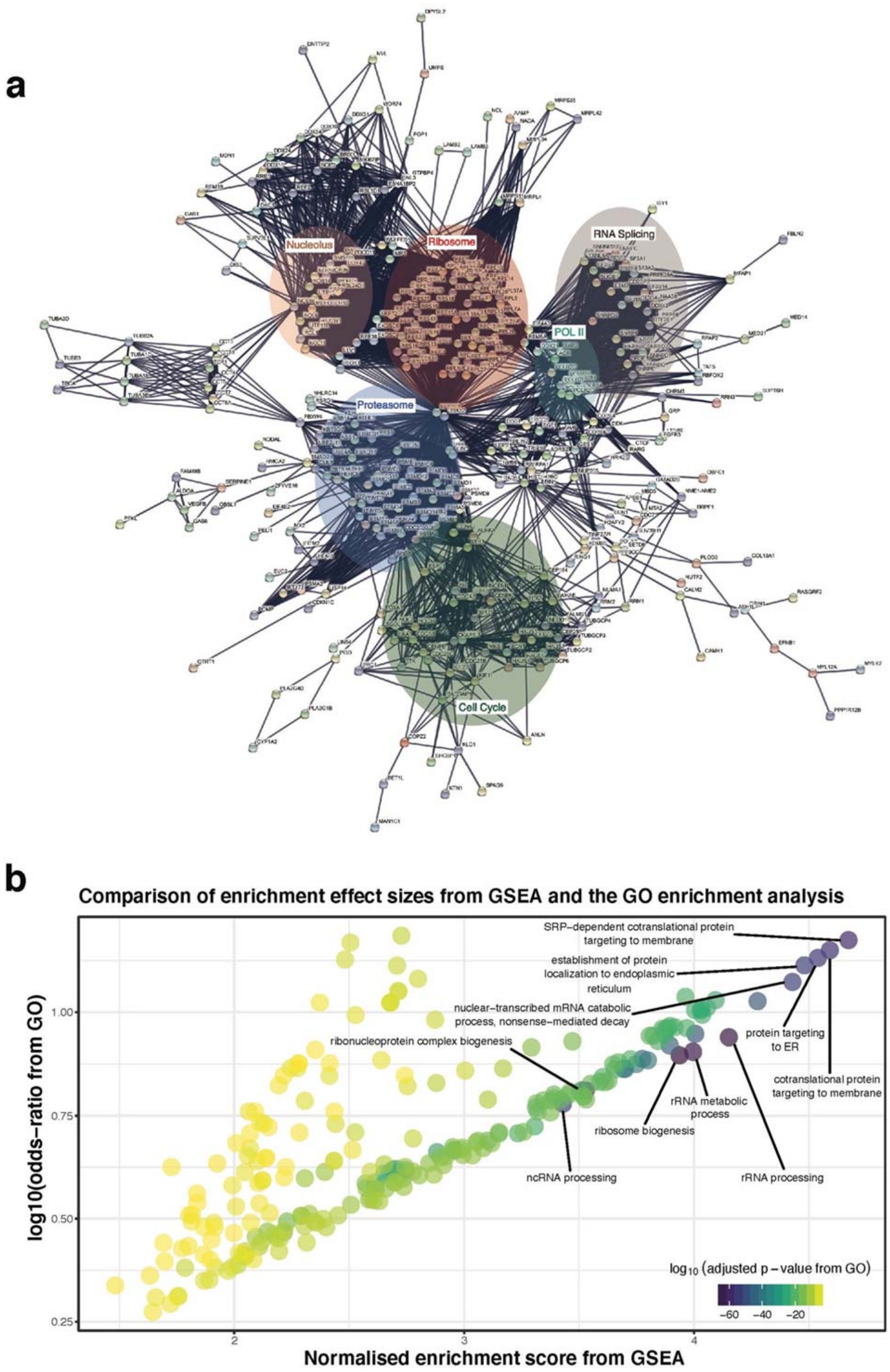


**Supplementary Figure 2: Further global analysis of the genome-wide high-throughput screen to identify components modulating p53 stabilisation.** (**a**) A fully annotated version of the STRING-Cytoscape analysis of candidates identified from the genome-wide screen (depicted in Fig. 1c of the main text). A comparison of the candidates from the screen using Gene Ontology (GO) and Gene Set Enrichment Analysis (GSEA, outlined in Methods), demonstrating that the core gene groups/ontologies correlate between both approaches (**b**).

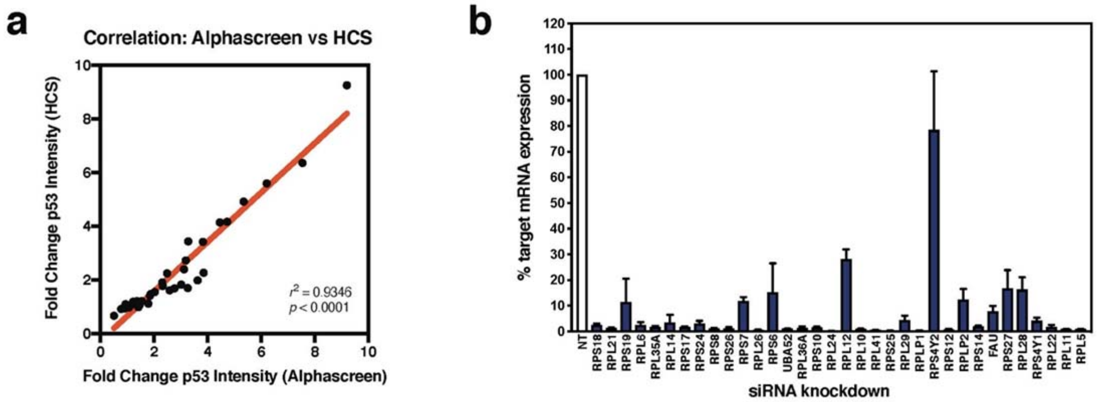


**Supplementary Figure 3: Additional analysis of ribosomal proteins identified from the screens.** We compared the results from the high-content screen (HCS) and the Alphascreen p53 assay approaches for the RPs screened; we observed a significant correlation ($r^2$ = 0.9346, $p$ < 0.0001) between the two methods (**a**). qRT-PCR analysis was used to confirm RP gene knockdown in A549 cells 24 hours post transfection with siRNA (n=2 or more biological experiments) (**b**); each target is normalised to cells transfected with non-targeting (NT) siRNA and depicted as a % expression relative to NT.

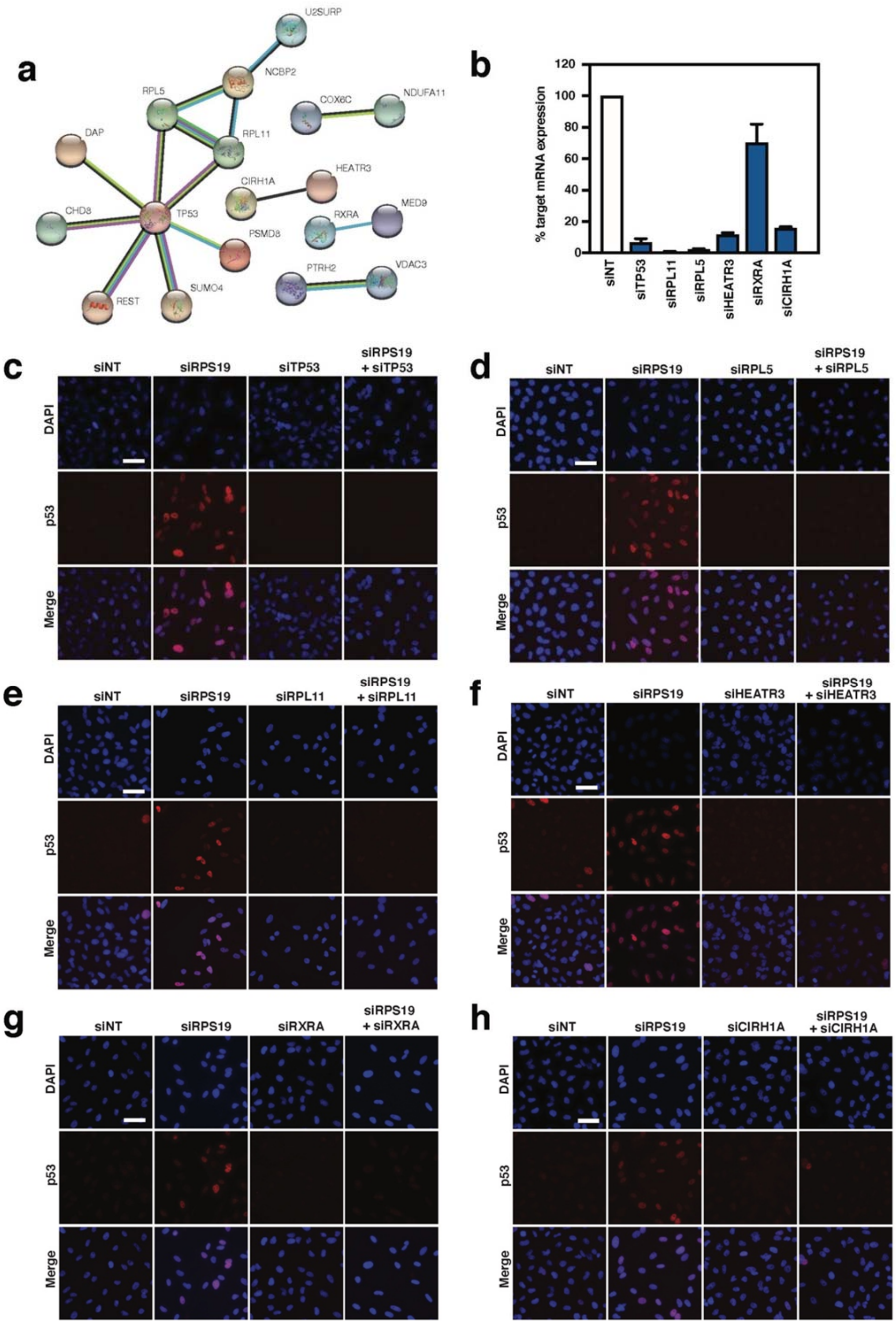


**Supplementary Figure 4 cont…**

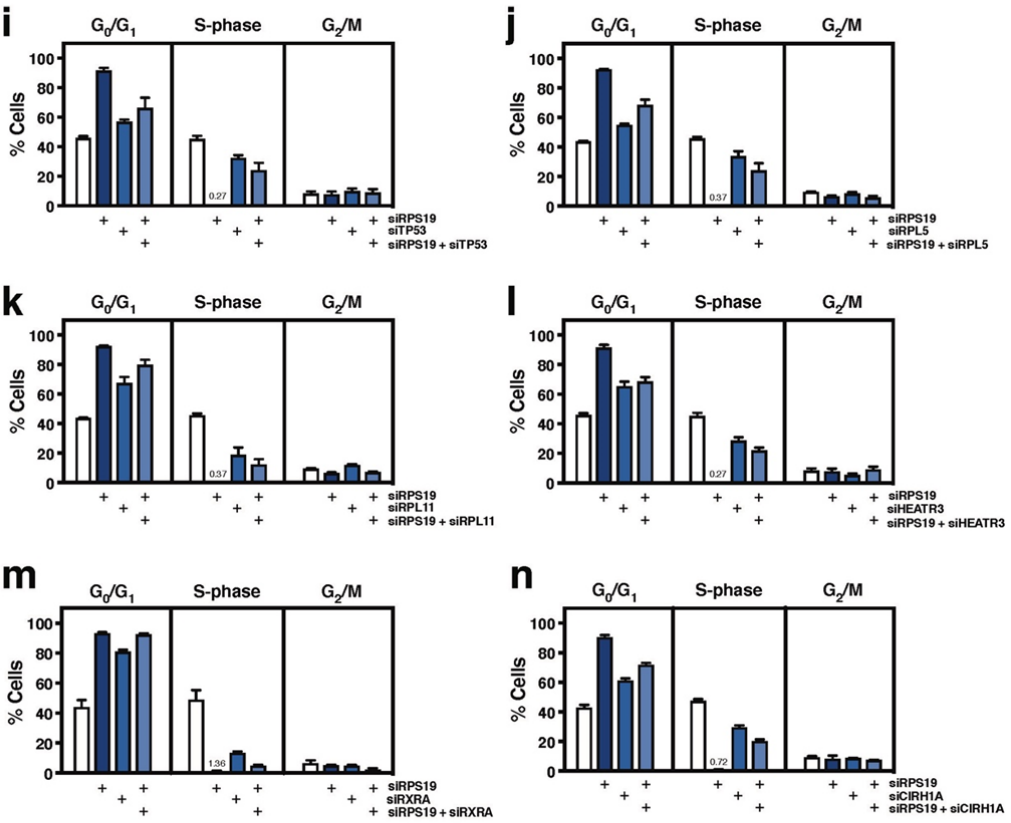


**Supplementary Figure 4: Analysis and validation of candidates identified from the high-throughput screen to identify genetic modulators of the canonical nucleolar surveillance pathway.** Analysis of the top candidates (64) from the "modifiers of ribosomal stress" screen using STRING analysis (only depicting those nodes with high confidence interactions) revealed a small subset of candidates interact directly with p53 (**a**). Further evaluation of expression of candidates identified in the screen (*TP53*, *RPL11*, *RPL5*, *HEATR3*, *RXRA* and *CIRH1A*) by qRT-PCR revealed target specific knockdown compared to non-targeting siRNA (siNT) 24 hours post-transfection in A549 cells (**b**). Data is graphed as mean +/- SD, n=2-3. Immunofluorescence evaluation of nuclear p53 protein expression in A549 cells transfected for 72 hours with RPS19 siRNA and co-transfected with siRNAs targeting TP53 (**c**), RPL5 (**d**), RPL11 (**e**), HEATR3 (**f**), RXRA (**g**) and CIRH1A (**h**) expression. Representative images captured at 20X magnification, scale bar = 50 µm, n=3 per candidate. Cell cycle analysis after 72-hour co-transfection with siRPS19 and siTP53 (**i**), siRPL5 (**j**), siRPL11 (**k**), siHEATR3 (**l**), siRXRA (**m**) or CIRH1A (**n**) in either $G_0/G_1$, S-phase or $G_2/M$. Data is graphed as mean +/- SD, n=2-3.

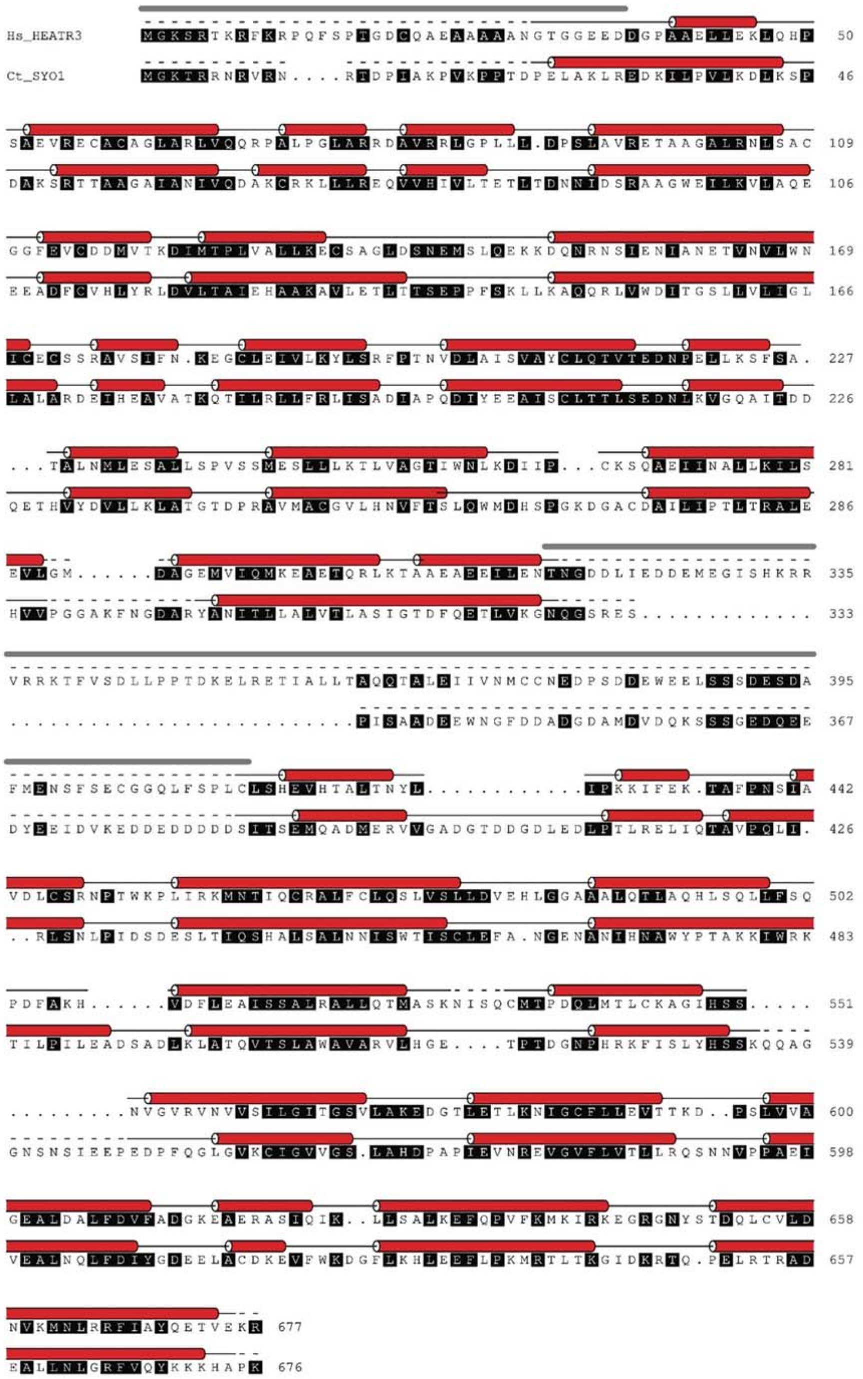


**Supplementary Figure 5: Secondary structure prediction-based sequence alignment of *Homo sapiens* (Hs) HEATR3 with *Chaetomium thermophilum* (Ct) Syo1.** Residues of the same similarity group are indicated in black according to the ALINE implementation[19]. Helical secondary structure of Syo1 (red tube, based on PDB 4GMO/5AFF) is indicated. HEATR3 secondary structure predictions (red tube) and predicted disordered regions (grey line) are indicated above the sequence. Sequence insertions are displayed as black dots, disordered regions in the crystal structure as a dashed line.

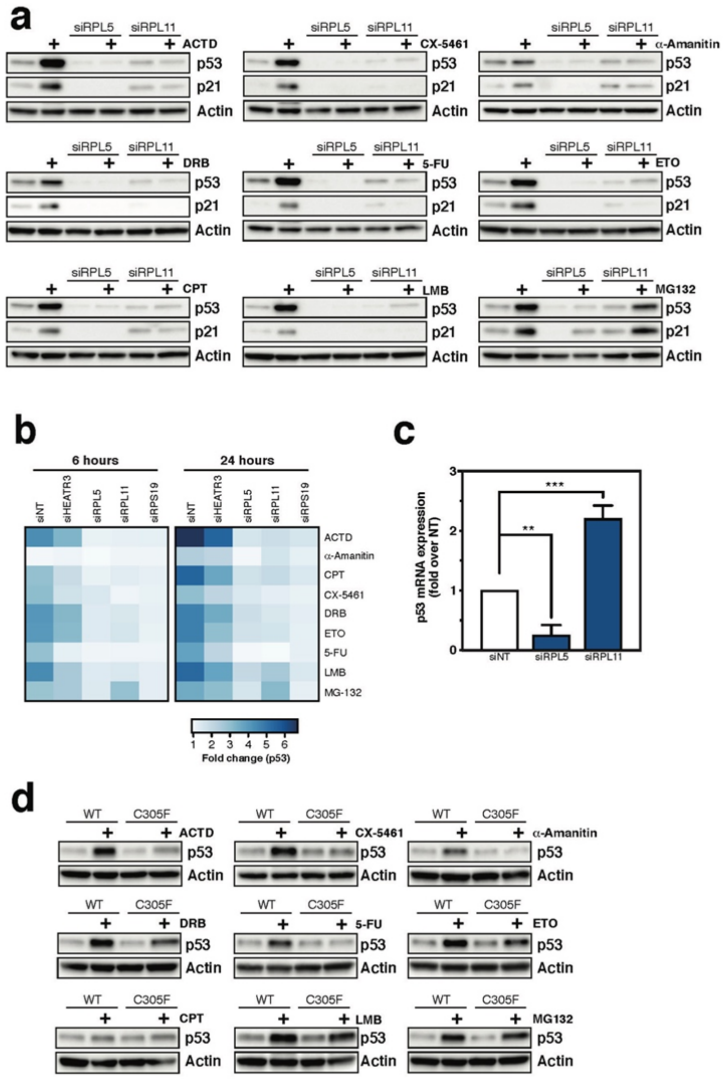


**Supplementary Figure 6: Further evaluation of p53 response to a broad range of genetic, pharmacological and pathophysiologic stresses when the NSP response is impaired.** We screened a panel of pharmacological agents and pathophysiological stressors when A549 cells were depleted of RPL5 and RPL11 for 48 hours. Cells were treated for 6 hours with pharmacological agents Actinomycin D (ACTD, 5 nM), CX-5461 (1µM), α-Amanitin (2.5 µM), Doxorubicin (DRB, 500 nM), 5-Fluoruracil (5-FU, 50 µM), Etoposide (ETO, 50 µM), Camptothecin (CPT, 50 nM), Leptomycin B (LMB, 10 ng/mL) or MG132 (10 µM) (note 24-hour data is presented in the main text). After treatment, protein was harvested and subjected to western blot analysis for p53 and p21 protein expression (**a**). Similarly, we utilised a high-content imaging approach to measure the effect of the aforementioned pharmacological agents on p53 stabilisation when HEATR3, RPL5, RPL11 and RPS19 were depleted from A549 cells for 48 hours and then treated for either 6 or 24 hours with each respective agent (**b**). In this instance, the data from the treatment with the agent was normalised to the vehicle-treatment for each individual siRNA condition (thus the heatmap presents the p53 differential for each agent within each individual siRNA condition). Quantitation of p53 mRNA (from equal cell number) in A549 cells transfected with RPL5 or RPL11 siRNAs for 72 hours and normalised to siNT-transfected cells (**c**, mean -/+ SD, one-way ANOVA with Dunnett's multiple comparisons test, $**p < 0.01$, $***p < 0.001$, n=3 biological experiments). The aforementioned pharmacological agent panel described above was also tested in mouse embryonic fibroblasts (MEFs) isolated from either Mdm2 wild-type (WT) or mice homozygous for the Mdm2 C305F mutation (C305F) for 6 hours to determine p53 expression (**d**, note 24-hour treatment panel is in Fig. 5C). Western blot images are representative of n=3-4 biological replicates.

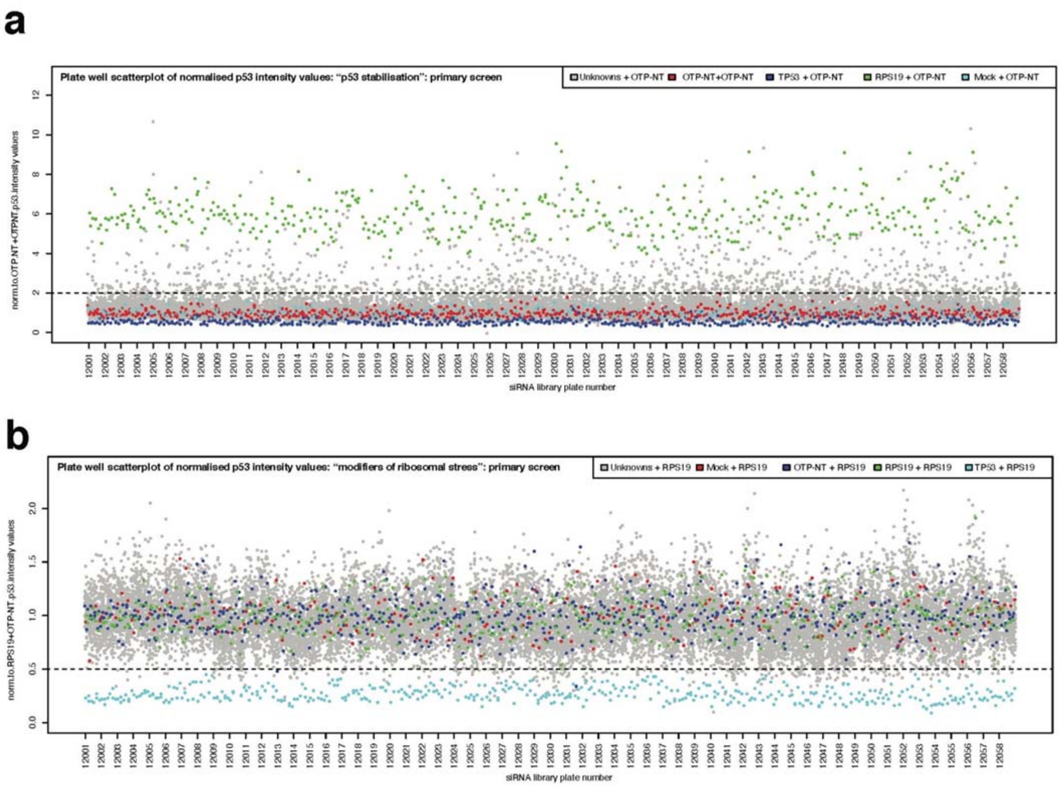


**Supplementary Figure 7: Scatterplots of normalised p53 intensity values from the primary siRNA screens.** We screened a total of 58 siRNA library plates in 384-well format (numbered 12001-12058) for each primary screen. The data was normalised to the specific control combination for each screen (OTP-NT+OTP-NT siRNA for the "p53 stabilisation screen", **a**, or to RPS19+OTP-NT siRNA for the "modifiers of ribosomal stress", **b**). Scatterplots of the individual normalised data points were then plotted to demonstrate the various screening controls (positive and negative) in relation to the unknowns; this was to ascertain over the course of the screen how the controls and unknowns were performing (as well as conducting Z' calculations, as outlined in the Methods). The dotted black lines represent the screening cut-off used for further analysis of screening candidates.

#### Supplementary File Descriptions

**Supplementary File 1:** Screening dataset for modulators of p53 stabilisation ('p53 stabilisation' full dataset – XLS file)

**Supplementary File 2:** STRING analysis for modulators of p53 stabilisation (used for Cytoscape analysis, XLS file)

**Supplementary File 3:** Gene Ontology analysis for modulators of p53 stabilisation (XLS file)

**Supplementary File 4:** LOCATE analysis for modulators of p53 stabilisation (XLS file)

**Supplementary File 5:** Ribosomal protein assembly analysis (XLS file)

**Supplementary File 6:** Screening dataset for genetic modulators of the canonical NSP ('modifiers of ribosomal stress screen' full dataset; XLS file)

**Supplementary File 7:** Data for heatmaps presented in Fig. 5a&b (XLS file)

**Supplementary File 8:** High-content screening acquisition and imaging analysis pipelines (PDF)

#### Supplementary References


1 Al-Hakim, A. K., Bashkurov, M., Gingras, A. C., Durocher, D. & Pelletier, L. Interaction proteomics identify NEURL4 and the HECT E3 ligase HERC2 as novel modulators of centrosome architecture. *Mol Cell Proteomics* **11**, M111 014233, doi:10.1074/mcp.M111.014233 (2012).

2 Durkin, M. E., Qian, X., Popescu, N. C. & Lowy, D. R. Isolation of Mouse Embryo Fibroblasts. *Bio Protoc* **3**, doi:10.21769/bioprotoc.908 (2013).

3 Birmingham, A. *et al.* Statistical methods for analysis of high-throughput RNA interference screens. *Nat Methods* **6**, 569-575, doi:10.1038/nmeth.1351 (2009).

4 Sloan, K. E., Bohnsack, M. T. & Watkins, N. J. The 5S RNP couples p53 homeostasis to ribosome biogenesis and nucleolar stress. *Cell Rep* **5**, 237-247, doi:10.1016/j.celrep.2013.08.049 (2013).

5 George, A. J. *et al.* A functional siRNA screen identifies genes modulating angiotensin II-mediated EGFR transactivation. *J Cell Sci* **126**, 5377-5390, doi:10.1242/jcs.128280 (2013).

6 Livak, K. J. & Schmittgen, T. D. Analysis of relative gene expression data using real-time quantitative PCR and the 2(-Delta Delta C(T)) Method. *Methods* **25**, 402-408, doi:10.1006/meth.2001.1262 (2001).

7 Trapnell, C., Pachter, L. & Salzberg, S. L. TopHat: discovering splice junctions with RNA-Seq. *Bioinformatics* **25**, 1105-1111, doi:10.1093/bioinformatics/btp120 (2009).

8 Anders, S., Pyl, P. T. & Huber, W. HTSeq--a Python framework to work with high-throughput sequencing data. *Bioinformatics* **31**, 166-169, doi:10.1093/bioinformatics/btu638 (2015).

9 Anders, S. & Huber, W. Differential expression analysis for sequence count data. *Genome Biol* **11**, R106, doi:10.1186/gb-2010-11-10-r106 (2010).

10 Mortazavi, A., Williams, B. A., McCue, K., Schaeffer, L. & Wold, B. Mapping and quantifying mammalian transcriptomes by RNA-Seq. *Nat Methods* **5**, 621-628, doi:10.1038/nmeth.1226 (2008).

11 Rueden, C. T. *et al.* ImageJ2: ImageJ for the next generation of scientific image data. *BMC Bioinformatics* **18**, 529, doi:10.1186/s12859-017-1934-z (2017).

12 Schneider, C. A., Rasband, W. S. & Eliceiri, K. W. NIH Image to ImageJ: 25 years of image analysis. *Nat Methods* **9**, 671-675, doi:10.1038/nmeth.2089 (2012).

13 de la Cruz, J., Karbstein, K. & Woolford, J. L., Jr. Functions of ribosomal proteins in assembly of eukaryotic ribosomes in vivo. *Annu Rev Biochem* **84**, 93-129, doi:10.1146/annurev-biochem-060614-033917 (2015).

14 Khatter, H., Myasnikov, A. G., Natchiar, S. K. & Klaholz, B. P. Structure of the human 80S ribosome. *Nature* **520**, 640-645, doi:10.1038/nature14427 (2015).

15 Chan, J. C. *et al.* AKT promotes rRNA synthesis and cooperates with c-MYC to stimulate ribosome biogenesis in cancer. *Sci Signal* **4**, ra56, doi:10.1126/scisignal.2001754 (2011).

16 Soding, J. Protein homology detection by HMM-HMM comparison. *Bioinformatics* **21**, 951-960, doi:10.1093/bioinformatics/bti125 (2005).

17 Zimmermann, L. *et al.* A Completely Reimplemented MPI Bioinformatics Toolkit with a New HHpred Server at its Core. *J Mol Biol* **430**, 2237-2243, doi:10.1016/j.jmb.2017.12.007 (2018).

18 Buchan, D. W. A. & Jones, D. T. The PSIPRED Protein Analysis Workbench: 20 years on. *Nucleic Acids Res* **47**, W402-W407, doi:10.1093/nar/gkz297 (2019).

19 Bond, C. S. & Schuttelkopf, A. W. ALINE: a WYSIWYG protein-sequence alignment editor for publication-quality alignments. *Acta Crystallogr D Biol Crystallogr* **65**, 510-512, doi:10.1107/S0907444909007835 (2009).

20 Jones, D. T. & Cozzetto, D. DISOPRED3: precise disordered region predictions with annotated protein-binding activity. *Bioinformatics* **31**, 857-863, doi:10.1093/bioinformatics/btu744 (2015).

21 Webb, B. & Sali, A. Comparative Protein Structure Modeling Using MODELLER. *Curr Protoc Bioinformatics* **54**, 5 6 1-5 6 37, doi:10.1002/cpbi.3 (2016).

22 Kressler, D. *et al.* Synchronizing nuclear import of ribosomal proteins with ribosome assembly. *Science* **338**, 666-671, doi:10.1126/science.1226960 (2012).

23 Franceschini, A. *et al.* STRING v9.1: protein-protein interaction networks, with increased coverage and integration. *Nucleic Acids Res* **41**, D808-815, doi:10.1093/nar/gks1094 (2013).

24 Szklarczyk, D. *et al.* STRING v11: protein-protein association networks with increased coverage, supporting functional discovery in genome-wide experimental datasets. *Nucleic Acids Res* **47**, D607-D613, doi:10.1093/nar/gky1131 (2019).

25 Shannon, P. *et al.* Cytoscape: a software environment for integrated models of biomolecular interaction networks. *Genome Res* **13**, 2498-2504, doi:10.1101/gr.1239303 (2003).

26 Yu, G., Wang, L. G., Han, Y. & He, Q. Y. clusterProfiler: an R package for comparing biological themes among gene clusters. *OMICS* **16**, 284-287, doi:10.1089/omi.2011.0118 (2012).

27 R Core Team (2018) R: a language and environment for statistical computing. (R Foundation for Statistical Computing, Vienna, Austria. http://www.R-project.org/).

28 Wang, J. Z., Du, Z., Payattakool, R., Yu, P. S. & Chen, C. F. A new method to measure the semantic similarity of GO terms. *Bioinformatics* **23**, 1274-1281, doi:10.1093/bioinformatics/btm087 (2007).

29 Subramanian, A. *et al.* Gene set enrichment analysis: a knowledge-based approach for interpreting genome-wide expression profiles. *Proc Natl Acad Sci U S A* **102**, 15545-15550, doi:10.1073/pnas.0506580102 (2005).

30 Sprenger, J. *et al.* LOCATE: a mammalian protein subcellular localization database. *Nucleic Acids Res* **36**, D230-233, doi:10.1093/nar/gkm950 (2008).
